# Redirecting cytomegalovirus immunity against pancreas cancer for immunotherapy

**DOI:** 10.1101/2025.05.15.654308

**Authors:** R Marrocco, J Patel, R Medari, P Salu, Meza E Lucero, S Brunel, A Martsinkovskiy, S Sun, K Gulay, M Jaljuli, E Mose, A.M Lowy, CA Benedict, T Hurtado de Mendoza

## Abstract

Immunotherapy shows limited success in pancreatic cancer, largely due to a low mutational burden and immunosuppressive microenvironment. Here we hypothesized that pre-existing antiviral immunity can be redirected to control pancreatic tumors. Cytomegalovirus (CMV, a β-herpesvirus) was chosen, as the majority of the population is infected and it induces an extremely large/broad memory T cell response. Mice latently infected with murine CMV (MCMV) were orthotopically implanted with pancreatic cancer cells and treated with systemic injections of MCMV T-cell epitopes. The therapy promoted preferential accumulation of MCMV-specific T cells within pancreatic tumors, delaying tumor growth and increasing survival. Immunophenotyping and scRNAseq analyses showed these T cells were highly activated and cytotoxic, leading to increased tumor necrosis and caspase-3 activation. Finally, therapy was enhanced when combined with subtherapeutic doses of gemcitabine chemotherapy. Together, these results show that CMV-specific T cells can be repurposed to combat pancreatic cancer.

**Significance:** Our studies reveal that CMV-specific viral memory T cells can be re-directed to control a solid tumor normally refractory to immunotherapy via a simple, intravenous injection of T cell peptide epitopes. This mutation agnostic approach has significant potential for the development of “off-the-shelf” therapeutics by stimulating pre-existing antiviral memory and it is widely applicable due to the high prevalence of CMV.

## Introduction

Immunotherapy has been most successful in the treatment of tumors with high mutational burdens such as melanoma and non-small cell lung cancer (1–4). These mutations result in the generation of neoantigens that the immune system recognizes as “non-self”, eliciting an anti-tumor immune response that can often be enhanced with immune checkpoint therapy. Other common tumors such as pancreatic cancer have a much lower mutational burden (5–7) coupled with an immunosuppressive tumor microenvironment (TME)(8, 9). Consequently, numerous strategies to enhance anti-tumor immunity through checkpoint blockade and/or by modulating the TME have been largely unsuccessful (10–12). Personalized therapies where tumor biopsies are sequenced for the emergence of potential neoantigens followed by vaccination are currently being tested. However, neoantigen prediction and subsequent therapy is not trivial, and this approach is currently very expensive and time consuming, oftentimes making it prohibitive.

A study of T cell antigens from long term survivors of pancreatic cancer showed that patients encoded neoantigens that mimicked pathogen sequences, suggesting they may be able to initiate an anti-tumor immune response under certain contexts (13). Here we hypothesized that delivering viral T cell epitopes to tumors might redirect this pre-existing antiviral immune memory and promote tumor clearance. Prior studies using subcutaneous tumor models by the Rosato and Cuburu groups *et al* (14, 15) showed that intra-tumoral injection of antiviral T cell peptide epitopes in mice previously infected with the virus led to growth arrest and even complete remission. However, direct intra-tumoral injection is not always feasible, and our approach to deliver epitopes systemically has the potential to be much more applicable to a variety of tumor types.

The tumor specificity of the iRGD peptide has been previously demonstrated by us and others (16–18), promoting systemic delivery of covalently linked or co-injected drugs to various solid tumors including breast, pancreatic, gastric, ovarian and others. iRGD interacts with αv integrins preferentially expressed by the tumor vasculature, inducing enhanced permeability through activation of the NRP-1 receptor and enhancing delivery of chemotherapy drugs, peptides, small molecules or even antibodies (19–25). Therefore, in this study we tested whether systemic delivery of CMV T cell epitopes co-administered with iRGD could induce an anti-tumor immune response.

We chose to repurpose cytomegalovirus (CMV) memory T cells because (i) >80% of the world’s population is infected, (ii) Immunodominant CMV-specific memory T cells show broad tissue residency and an effector memory phenotype in people and mice and (iii) ∼10% of all circulating CD4 and CD8 T cells are CMV-specific (26, 27). In the present study, mice were infected with MCMV until viral latency was established (>2 months), were orthotopically implanted with KPC pancreatic cancer cells and once tumors were palpable were injected with MHC class I and II MCMV peptide epitopes +/− iRGD (MCMVp therapy). Mice responded to treatment as evidenced by enhanced survival, delayed tumor growth and increased tumor apoptosis and T cell infiltration. Both immunophenotyping by flow cytometry and scRNAseq analyses of tumors showed that MCMV-specific TILs were highly activated and cytotoxic. Surprisingly, the tumor specific homing of the MCMV T cells and the antitumor effects of the MCMVp therapy were independent of iRGD. Together, these data show that mobilizing pre-existing viral memory T cells shows clinical efficacy against pancreatic tumors.

## Results

### MCMV memory T cells can be redirected to fight pancreatic tumors

In order to induce an MCMV memory T cell pool, we infected C57bl6/J (B6) mice at 4-5 weeks of age and waited a minimum of 2 months to allow the establishment of latency, defined by maintenance of the viral genome without detectable lytic replication. MCMV-infected and non-infected control mice were then surgically implanted with KPC1242 tumor cells in their pancreas. When tumors were palpable (∼ 9-11 days post tumor cell injection (dpi)), mice were treated bi-weekly with either vehicle (DMSO/PBS) or iRGD plus MCMVp therapy (three 15-mer CD4 T cell peptide epitopes derived from the viral proteins m09, m25 and m142 and three 8/9-mer CD8 T cell peptide epitopes derived from the viral proteins m38, m45 and IE3) (Fig. 1A). Tumor growth was monitored by ultrasound and mouse weights were taken twice a week.

**Figure 1.**
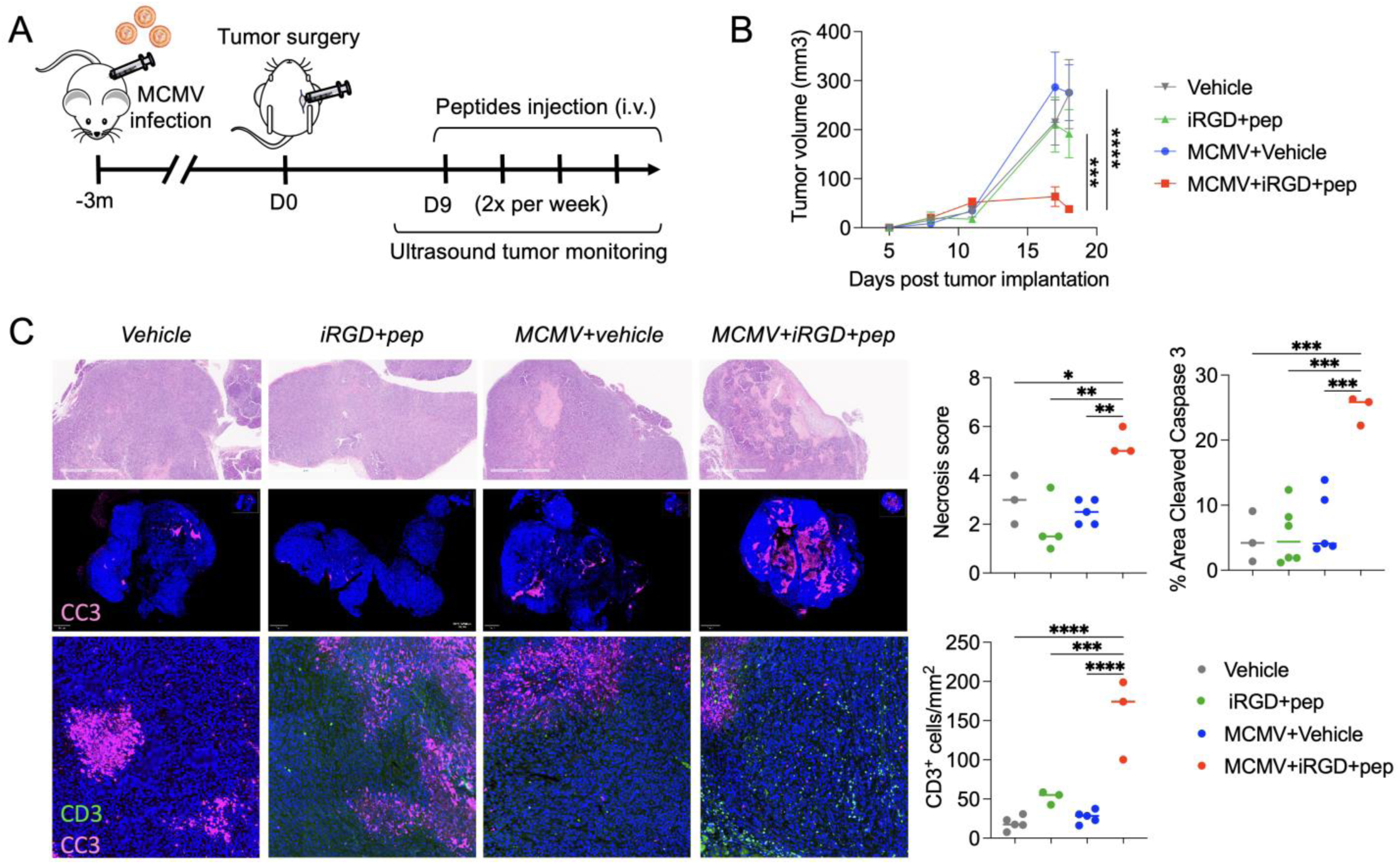
MCMV memory T cells can be redirected to fight pancreatic tumors. **A**, Protocol overview. Each treatment consists of an iRGD injection (300µg) followed by injection of 50µg of each MCMV peptide. **B,** KPC1242 tumor growth over time, monitored by ultrasound. (n=4-5/group). **C,** H&E and immunofluorescent analysis of the tumors at endpoint looking at apoptosis by cleaved caspase 3 (CC3-Red) and T cell infiltration by CD3 (green) staining. Necrosis score was attributed blindly by a histopathologist. Results were compared by one-way ANOVA with Tukey correction, and repeated measures by two-way A NOVA with Sidak correction. *P < 0.05; **P < 0.01; ***P < 0.001; ****P < 0.0001.

Infected mice treated with iRGD + MCMVp therapy showed a dramatically reduced tumor growth, while infection alone or treatment of uninfected mice showed no benefit (Fig. 1B and S1A bottom panel). Histological analysis of the tumor tissue showed increased necrosis, cleaved caspase-3 and CD3^+^ T cell infiltration (Fig. 1C). On the other hand, the same analysis in the liver showed increased T cell infiltration but no cleaved caspase 3 staining (Fig. S1D) Despite the beneficial effects of MCMVp therapy, some toxicity resulting in significant weight loss was observed in the infected and treated group, possibly due to the high doses of MCMVp used initially (50 µg of each) (Fig. S1A top panel). This was associated with a massive expansion of highly proliferative (Ki67) and activated (CD69) MCMV specific CD8 T cells in the tumor and spleen (>80% of total T cells), on the other hand, the expansion of MCMV CD4 T cells in the spleen was minimal (Fig. S1B and S1C). This prompted us to perform a dose titration of the MCMVp to see if a therapeutic window lacking systemic toxicity could be found. After testing different doses of CD4 and CD8 peptide epitopes (Fig. 2A), we determined that 1µg of CD8 peptides and 50 µg of CD4 peptides (Dose A) was optimal. Notably, this dose of MCMVp therapy still imposed tumor growth control without inducing measurable weight loss. A survival study using this dose showed a 68% increase in survival with a median of 25 days for infected mice treated with vehicle vs 42 days for infected mice that underwent MCMVp therapy (p=0.0027) (Fig. 2A). We next assessed the effect that injection of iRGD or MCMVp alone had on tumor growth and survival compared to the combination. As expected, iRGD alone had no beneficial effect. Surprisingly, MCMVp alone had the same effect on T cell infiltration, tumor growth and survival than its combination with iRGD (Fig. S2A-C). Interestingly, 3 immunodominant MCMV-specific T cell populations (m25 CD4 T, and m38/m45 CD8 T cells) were still preferentially enriched in the tumor compared to the spleen and liver despite the absence of iRGD, indicating an iRGD-independent mechanism favors their tumor localization (Fig. S2D). Given these results, the rest of our studies were conducted without addition of iRGD.

**Figure 2.**
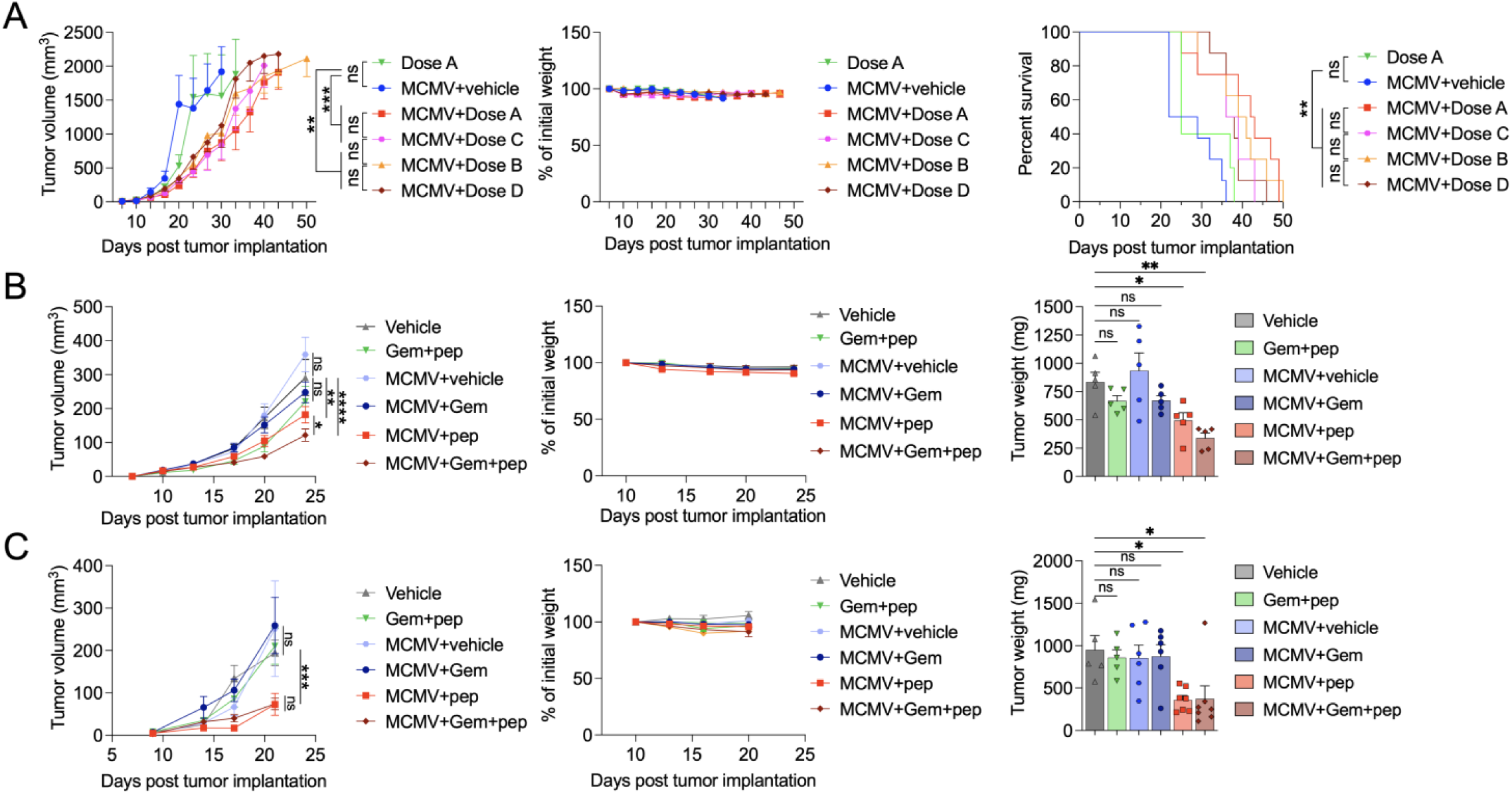
MCMVp therapy results in delayed tumor growth with increased survival and can be enhanced with chemotherapy. **A**, KPC1242 tumor growth curves using various doses of MCMVp therapy (Dose A 50µg CD4p/1µgCD8p, Dose B 50µg CD4p/0.1µgCD8p, Dose C 10µg CD4p/3µgCD8p, Dose D 10µg CD4p/1µgCD8p). Normalized mouse weight (middle) and Kaplan-Meier survival graph (right). (n=5-8/group). **B,** Dose A was used in combination or not with gemcitabine (Gem). Normalized mouse weight (middle) and tumor weight at endpoint (right). (n=5-10/group) **C,** B6×129 F1 hybrid mice were injected with the KPC46 tumor cell line and treated as in B. Normalized mouse weight (middle) and tumor weight at endpoint (right). (n=5-7/group). Results were compared by one-way ANOVA with Tukey correction, and repeated measures by two-way A NOVA with Sidak correction. *P < 0.05; **P < 0.01; ***P < 0.001; ****P < 0.0001.

Since high PD-1 expression was observed in MCMV-specific TILs (Fig. S1C), we assessed whether checkpoint blockade could further enhance MCMVp therapy-dependent tumor control. As KPC tumors contain high levels of IL-10 producing myeloid-derived suppressor cells (MDSC), we also combined our treatment with a blocking anti-IL-10R antibody. However, no additional benefits to MCMVp therapy were observed, even when both PD-1 and IL-10R blockade were combined (Fig. S2E). This was the last treatment study performed in combination with iRGD. We then sought to assess whether combining MCMVp therapy with a chemotherapeutic agent could show additional benefit. We decided to use Gemcitabine (Gem), one of the standard of care drugs, known for its potential to reduce the levels of MDSCs and enhance T cell immunity, especially when used at low doses (5mg/kg) (28–30). In this treatment study, we found that while Gem lacked any anti-tumor effect on its own at 5mg/kg, co-treatment with MCMVp showed significant benefit over either monotherapy, with no apparent toxicity, resulting in a further delay in tumor growth and significantly decreased tumor weights at end point (Fig. 2B).

In order to show that the benefits of MCMVp therapy were not restricted to a single pancreatic tumor model, we used the C57Bl6/J (B6) x129SvJ (129) F1 hybrid model implanted with the syngeneic KPC46 tumor cell line. Similar to KPC1242 tumors, neither MCMV infection without MCMVp therapy or peptide epitope treatment of uninfected mice impacted tumors. Only infected mice receiving MCMVp therapy showed a significant decrease in tumor growth (Fig. 2C). In this model, MCMVp therapy alone showed markedly enhanced efficacy compared to KPC1242 in B6 mice, with Gem having no additional benefit. Taken together, we have shown that harnessing pre-existing CMV memory T cell responses has major therapeutic benefits in two different aggressive, orthotopic pancreatic tumor models.

### MCMV T cells expand following treatment and preferentially localize to the tumor

We next sought to characterize the phenotype and effector function of MCMV specific T cells in spleen, liver and tumor in the various therapeutic regiments described above (Fig. 3 and Fig. S3). First, we determined the total number of CD8 T cells and CD4 Tconv (CD4T effector cells not including Treg) and saw (i) no differences in the spleen, (ii) increased CD8 and CD4 Tconv in the liver of infected mice receiving MCMVp therapy (+/−Gem) and (iii) increased CD8 T cells in tumors of the same 2 therapeutic groups. (Fig. S3A). To assure specificity, MCMV-specific T cells were identified using both MHC-I and -II tetramers labeled in both APC and PE, and only double-positive cells were analyzed (Fig. 3 A, D). MCMV specific CD4 Tconv cells were detected exclusively in tumors from infected mice receiving MCMVp therapy. The spleen contained a low proportion of MCMV T cells, while the liver and tumor had higher and more comparable levels. The dominant epitope in the liver was m142, while tumors were more enriched with m25 specific T cells (Fig 3A, B). Quantification of the absolute number of m09, m25 and m142 specific Tconv cells in tumors (Fig. 3C), livers and spleens (Fig. S3B) paralleled their proportions shown in Fig 3B. In addition, CD4^+^FoxP3^+^ MCMV-specific regulatory T cells were analyzed, and composed a much lower proportion than CD4 Tconv (Fig. S3D), suggesting that *in vivo* expansion of MCMV-specific CD4 T cells results in the differentiation of a bonified antiviral effector CD4 T cell rather than an immunoregulatory one.

**Figure 3.**
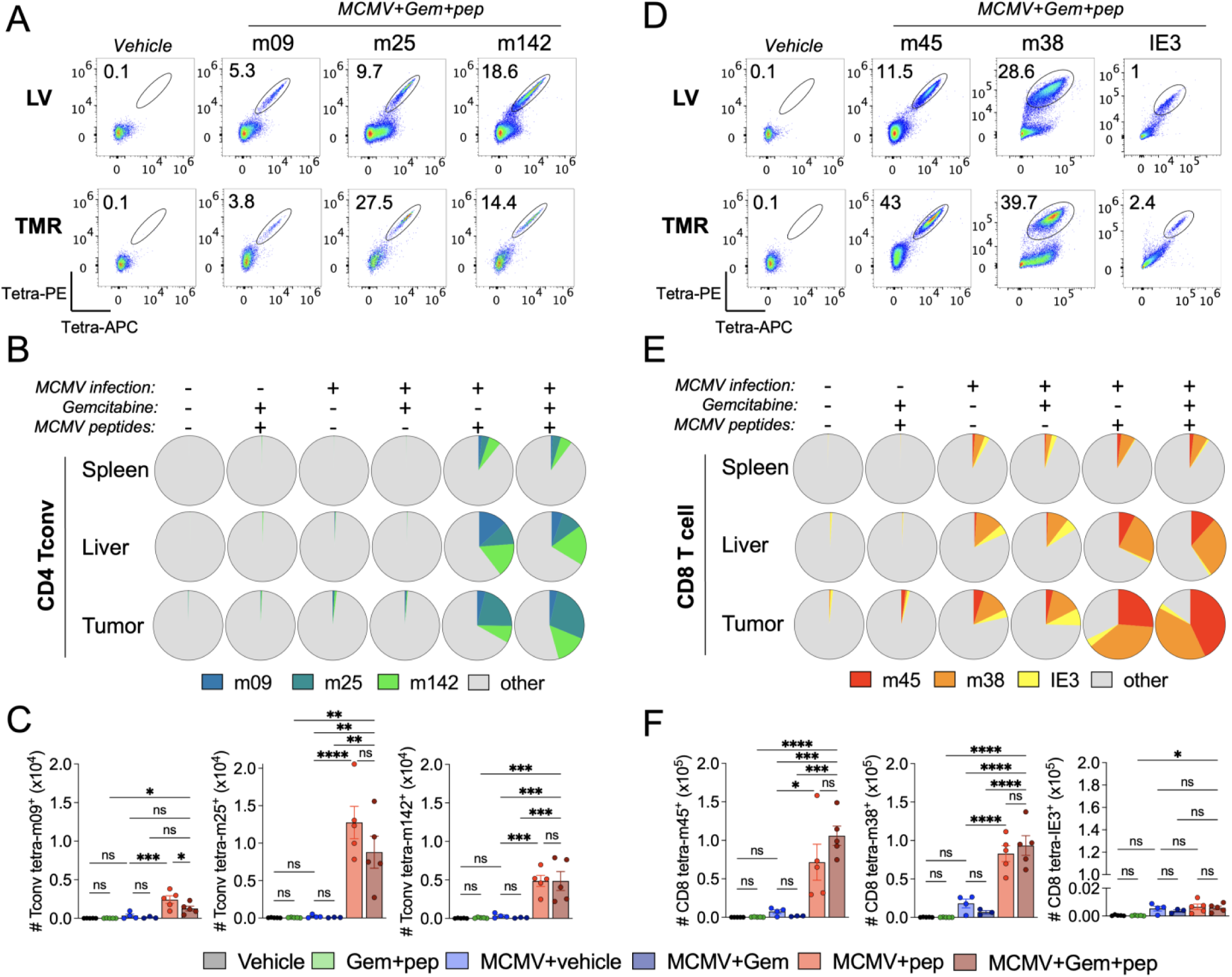
MCMV T cells expand after MCMVp therapy and preferentially localize to the tumor. Mice were implanted with KPC 1242 cells and treated as previously described in Fig 2B. At day 25 p.i. spleen, liver, and tumors were collected to analyze TILs by flow cytometry. (n=5/group). **A**, **D**, Representative dot-plot of tetramer binding MCMV-specific CD4 Tconv (A) and CD8 T cells (B). **B**, **E**, Mean proportions of MCMV tetramer binding T cells among total CD4 Tconv (B) and CD8 T cells (E). **C**, **F**, Absolute number of tumor-resident MCMV tetramer-binding CD4 Tconv (C) and CD8 T cells (F). Results were compared by one-way ANOVA with Tukey correction. *P < 0.05; **P < 0.01; ***P < 0.001; ****P < 0.0001.

Notably, while MCMV-specific T cells were detectable in tumors from infected mice in the absence of MCMVp therapy (Fig. 3E) it did not translate to a significantly higher absolute number of these T cells in the tumor (Fig. 3F), due to low overall T cell infiltration. After MCMVp therapy we observed an expansion of m38 and m45 CD8 T cell proportions in the liver as well as the tumor (Fig. 3D, E), commensurate with an increase in their absolute numbers in tumors (Fig 3F) and livers (Fig.S3C), while IE3 CD8 T cells failed to significantly expand and infiltrate the tumor. Taken together, these data show that MCMVp therapy in infected mice selectively enhanced the presence of m25, m38 and m45 specific T cells in the tumor and to a lesser extent in the liver. Despite the increase in MCMV specific T cells in the liver following MCMV therapy, histological analysis by cleaved caspase 3 staining showed no tissue damage, even at the highest doses of MCMVp (Fig S1D).

### MCMV-specific CD8 T cells show a marked effector phenotype

The pancreatic cancer TME is highly immunosuppressive, tending to inhibit T cell effector function and promote tumor growth. Consequently, we analyzed the phenotype and function of tumor-resident MCMV-specific CD8 T cells to assess their potential anti-cancer activity. The expression levels of activation markers (CD44, CD69), costimulatory receptors (CD226), inhibitory receptors (PD1, LAG3, and TIM3) and cytotoxic molecules (Prf, GzmA, and GzmB) were assessed. All of these markers except GzmA were significantly increased in m45-specific CD8 TILs from infected mice receiving MCMVp therapy (Fig. 4A). Of note, GzmB levels were significantly higher in mice treated with MCMVp+Gem, providing a possible mechanistic explanation for the enhanced tumor control in this group compared to MCMVp therapy alone. M45-specific CD8 T cells also showed a very similar phenotype in the liver, suggesting a conserved phenotype in multiple tissues (Fig. S4A). While PD1, LAG3 and TIM3 are often called “exhaustion markers”, they don’t define functional exhaustion as they are also upregulated early during T cell activation to dampen excessive inflammation (31). Consistently, we observed strong expression of activation markers and cytotoxic molecules in T cells expressing these immune checkpoint molecules, arguing against an exhausted state. *In vitro* re-stimulation of the tumor and liver cells with a mix of the 6 MCMV peptides used for treatment resulted in robust IFNγ and TNF production by CD8 T cells, further indicating these cells were not exhausted (Fig. 4B, S4D). Of note, MCMV-specific T cells also established tumor residence in infected mice that did not receive MCMVp therapy, as shown by tetramer binding (see Fig 3E) and cytokine production (Fig 4B). MCMVp therapy did further increase both IFNγ and TNF production by these TILs. This enhanced IFNγ was most evident in the MCMVp+Gem group, correlating with greater tumor growth control, and the same was true for IFNγ production by CD4 Tconv in MCMVp+Gem treated mice (Fig. 4B).

**Figure 4.**
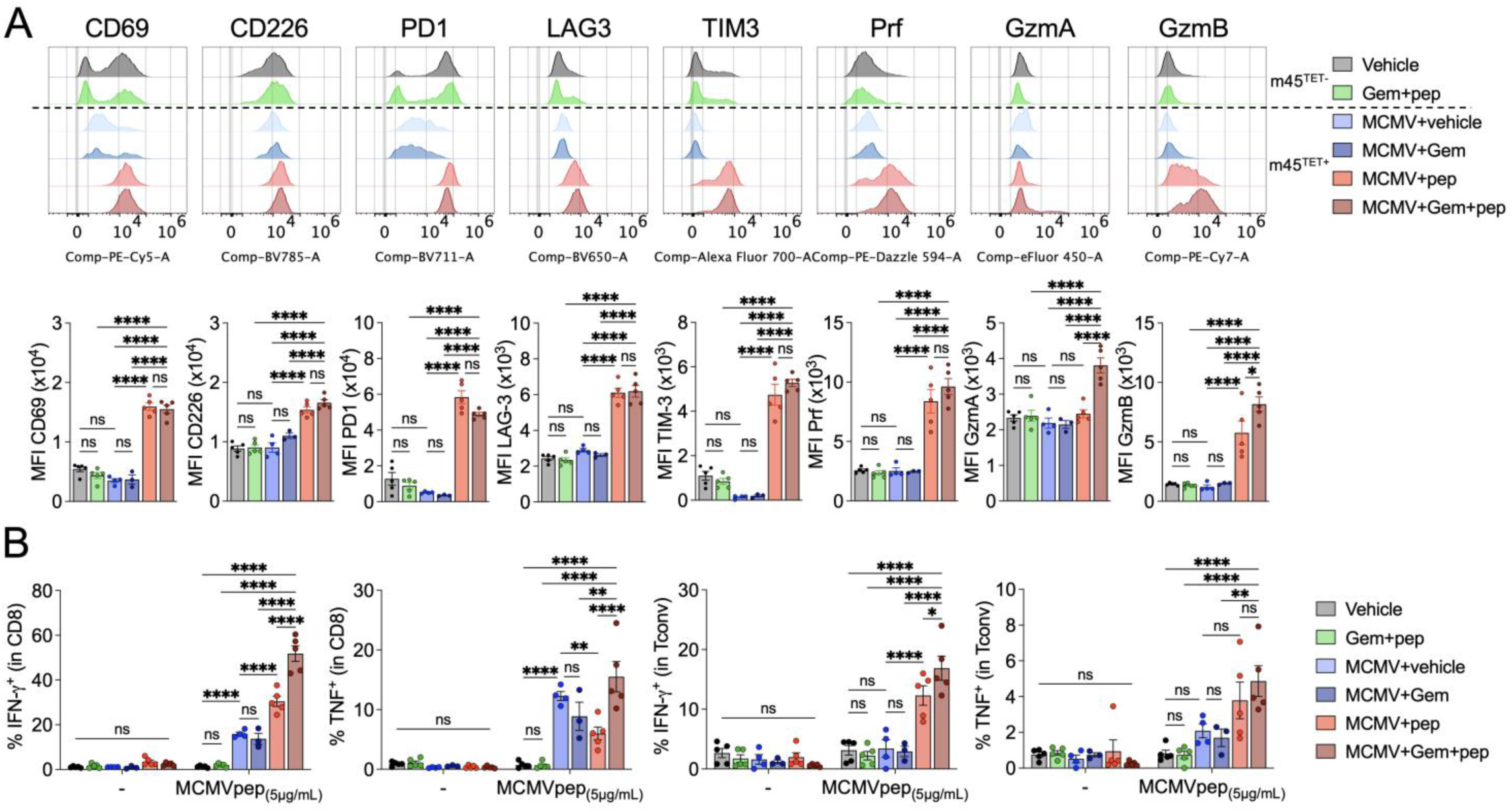
MCMV T cells are highly activated and cytotoxic within tumors. Immunophenotyping and cytokine production analysis of the TILs described in Fig 3. (n=5/group). **A**, Representative histograms of m45-specific CD8 T cells isolated from tumors (top) and mean fluorescence intensity (MFI) quantification (bottom). **B**, Tumors were subjected to CD45+ TIL magnetic separation and the purified cells were incubated with the 6 MCMV peptides for 4h, in the presence of GolgiPlug. The graphs show IFNγ and TNF production in CD8 or CD4 T conv cells. Results were compared by one-way ANOVA with Tukey correction. *P < 0.05; **P < 0.01; ***P < 0.001; ****P < 0.0001.

Following MCMV infection, m45-specific CD8 T cells show conventional early expansion/contraction kinetics and display a central memory T cell phenotype. Conversely, m38-specific CD8 T cells display unique kinetics. They undergo early expansion paralleling m45, but only weakly contract and continue to “inflate” in numbers over time displaying an effector-memory T cell phenotype. Despite the intrinsic differences in these two distinct antiviral CD8 T cell populations, the phenotype of m38 and m45 populations was highly similar in both liver and tumor (Fig. 4A, S4A-C). GzmA and GzmB expression was significantly higher in liver-resident m38 compared to m45 T cells in infected mice not receiving MCMVp therapy (Fig. S4A and B). However, once treated with MCMVp ± Gem, both CD8 T cell populations showed similar Gzm expression in the liver. In the tumor, both these CD8 T cell populations expressed lower GzmA than in the liver. Surprisingly, tumor resident m38 T cells downregulated GzmA expression after MCMVp therapy, but upregulated GzmB (Fig. S4C). This downregulation of GzmA may result from the TME, but, importantly, did not prevent the beneficial effect of MCMVp therapy.

### MCMVp treatment induces profound changes in the tumor microenvironment

To assess the effect of MCMVp therapy on the TME, we performed scRNAseq of whole tumors from MCMV infected mice treated with vehicle or MCMVp (Fig. 5A). The major tumor cell populations were epithelial cells, macrophages, T cells, fibroblasts, and dendritic cells. As shown (Fig. 5A, B, Fig S5A), the most significant change in the MCMVp therapy group was a marked increase in T cells (0.27%Veh vs 5.7%MCMVp). Additionally, a trend was seen for decreased pancreatic tumor cells (80%Veh vs 66%MCMVp), and increased proportion of macrophages (17%Veh vs 25%MCMVp) and dendritic cells (0.7%Veh vs 1.87%MCMVp). When lymphocytes were subclustered into B cells, CD4/CD8 T cells and NK cells we observed that the proportion of CD8 T cells was markedly increased following MCMVp therapy, while B and NK cells remained largely stable (Fig. S5B). GSEA analysis of epithelial cells revealed an increase in type-I and type-II IFN response genes and a decreased expression of genes involved in angiogenesis, KRAS signaling and epithelial-mesenchymal transition (EMT) (Fig. 5C). Volcano plots showed a notable increase in antigen presentation genes such as H2-D1, H2-K1 and B2m, consistent with an enhanced anti-tumor immune response. On the other hand, EMT-related genes like Acta2 and genes associated with tumor progression such as Megf10 or pleiotrophin (ptn) were significantly downregulated (Fig. S5C Left panel).

**Figure 5.**
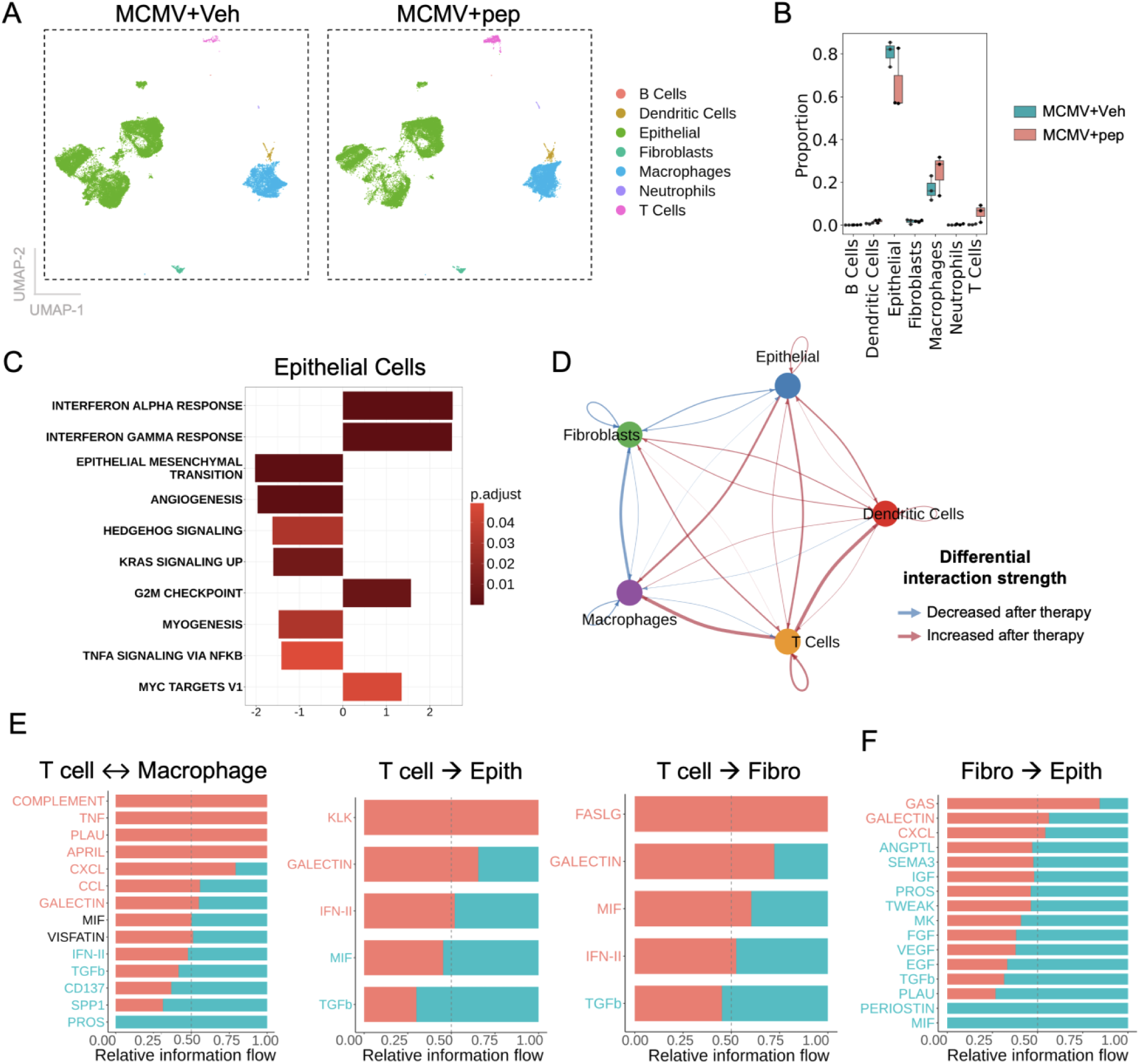
MCMVp treatment induces profound changes in the tumor microenvironment. Infected mice bearing tumors were treated with vehicle or MCMVp as previously described, tumors were collected and used for 5’scRNAseq (n=3/group, each sample being the pool of 1-2 mice). **A**, UMAP distribution. **B**, Boxplot of cellular proportions in the tumor. **C**, GSEA analysis of epithelial cell DEGs. **D** CellChat analysis of potential cell-cell interactions in the tumor. Arrowheads indicate the directionality of the interactions. **E, F**, Relative quantification of the different pathways involved in the potential cell-cell interactions in the tumor shown in D. Double arrows indicate bi-directional interactions, simple arrows indicate unidirectional interactions.

Fibroblasts and other stromal cell populations are known to support tumor growth and shape the TME (32), in the context of MCMVp therapy we observed a significant increase in MHC class II genes, such as H2-Aa, H2-Ab1 and H2-Eb1 associated with CD4 T cell mediated anti-tumor immune responses (Fig.S5C Middle panel). In addition, we looked at DEGs in macrophages and found an upregulation of Stat1 and Irf1, two IFN response related genes, and Tap1, involved in peptide processing, consistent with an enhanced immune response (Fig.S5C Right panel).

We then utilized CellChat to investigate how cell-cell interactions may be impacted by MCMVp therapy. This analysis suggested reduced interactions between fibroblasts and both epithelial cells and macrophages, while T cell interactions with macrophages, dendritic cells and epithelial cells were increased (Fig. 5D). This conclusion was based on the apparent increased expression of chemokines, complement, APRIL/*Tnfsf13*, and Tnf mediated crosstalk between T cell and macrophages. Decreased TGF-β and SPP1 dependent pathways was also observed (Fig 5E), which are involved in maintaining the immunosuppressive TME phenotype (33, 34). TGF-β dependent signaling networks are also decreased between T cells, epithelial cells and fibroblasts (Fig 5E, F). In turn, MCMVp therapy increased potential *FasLg* interactions between T cells and fibroblasts, but not between T cells and epithelial cells. Importantly, fibroblast-epithelial cell interactions involving growth factors (FGF, VEGF, EGF) were reduced following MCMVp therapy, which would be consistent with a better prognosis (Fig. 5F). Taken together, our results show that with a peptide-based therapy we are able to induce significant changes in the TME that lead to improved survival and better prognosis.

### MCMV therapy leads to tumor infiltration of activated and cytotoxic MCMV TCRαβ cells

To further explore the TIL phenotype following MCMVp therapy, and whether ‘bystander activation’ of T cells that are not virus specific may occur, a second scRNAseq on FACS sorted CD3^+^CD4^+^ and CD3^+^CD8^+^ T cells was performed (Fig. 6). ∼50,000 T cells were analyzed for both gene expression and V(D)J CDR3 sequences to assess the overall TCRαβ repertoire. Bioinformatic analysis yielded 20 distinct T cell clusters (Fig. 6A), with 13 composed by CD8 T cells, 5 representing CD4 Tconv and 2 FoxP3+ CD4 Treg (Fig. 6A, B).

**Figure 6.**
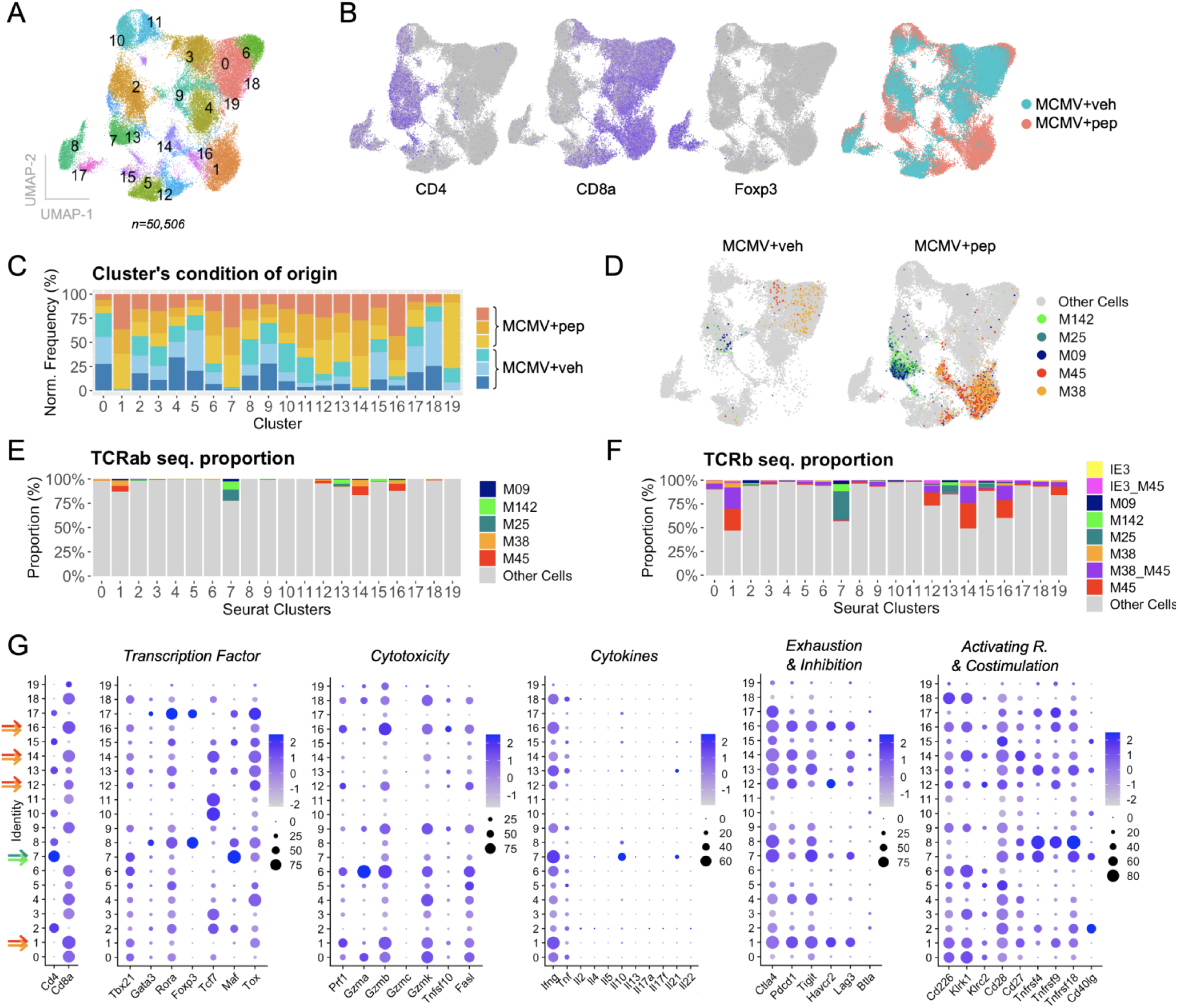
MCMV therapy leads to tumor infiltration of activated and cytotoxic MCMV TCRαβ cells. A different cohort of mice treated as in Fig 5, was used to purify T cells by FACS sorting for CD3, CD4 and CD8 (n=3/group, each sample being the pool of 1-2 mice). The purified T cells were used for 5’scRNAseq and TCR sequencing. **A**, T cell clustering and UMAP distribution. **B**, Feature plot mapping of CD4, CD8 and regulatory T cells within these clusters and overlay of the 2 conditions (Vehicle-Blue, MCMVp-Red)**. C**, Graph showing the frequency of each sample in each of the predefined clusters after normalizing the amount of cells per sample **D**, Dimplot representing the detection of specific MCMV-associated matched CDR3 sequence for TCRα and TCRβ. **E**, Proportion of MCMV-specific TCRαβ detected per cluster. **F**, Proportion of MCMV-specific TCRβ detected per cluster. **G**, Dotplots featuring selected gene-sets and expression levels among clusters. Arrows point to clusters containing high proportion of MCMV-specific T cells (red=m45, orange=m38, green=m142, blue=m25).

Analysis of T cells in mice receiving MCMVp therapy clearly showed the emergence of 5 new clusters (Fig. 6B, S6A). Clusters 1, 7, 12, 14 and 16 were largely exclusive to the MCMVp therapy group (Fig. 6C), with cluster 7 being CD4 Tconv and the other four composed of CD8 T cells. To assess whether these clusters contained MCMV-specific T cells, we analyzed their TCRα and TCRβ CDR3 sequences (Fig. 6D). We have previously determined the paired TCRαβ sequences from MCMV CD4 Tconv that arise during MCMV infection of B6 mice (manuscript in preparation), and others have published this for MCMV-specific CD8 T cells (35, 36). In the CD4 Tconv cluster 7, we observed a strong signal for m25 and m142-associated CDR3s, and an intermediate signal for m09. In cluster 7, ∼25% of CD4 T cells in the MCMVp group contain MCMV TCRαβ sequences compared to ∼1% in infected mice not receiving therapy. Similarly, we detected a large proportion of m45 and m38-associated CDR3s in clusters 1, 16 and 14, and a weaker signal in cluster 12 (Fig. 6D and E). Next, we repeated the analysis looking exclusively at TCRβ sequences, since more MCMV-specific TCRβ sequences are known, compared to TCRα sequences (Fig. 6F, S6B). Up to 50% of cluster 1 is composed of MCMV-specific CD8 T cells, half that express m45-specific CDR3, and the other half expressing CDR3 ascribed to both m45/m38 in the literature (35, 36) (Fig. 6F). Since cluster 1 is the second most abundant cluster (Fig. S6C), we can conclude that MCMV-specific CD8 T cells constitute the majority of the tumor infiltrating T cells.

We then analyzed all 20 clusters for the expression of key transcription factors, cytotoxic molecules, cytokines and co-signaling receptors that regulate T cell effector function (Fig. 6G). All clusters induced by MCMVp therapy expressed high levels of *Tbx21* (T-bet) and *Ifnγ*. Cluster 1 expressed very high levels of *Prf1, Gzmb*, *Gzmk* and *Fasl*, indicative of high cytotoxic potential. As seen by flow cytometry (see Fig 4 and S4), elevated expression of co-inhibitory receptors was observed in cluster 1 (*Pdcd1*, *Havcr2*, *Lag3*), as well as *Ctla4* and *Tigit*. However, enhanced levels of several co-stimulatory receptors were also seen (*Cd226*, *Klrk1*, *Cd27*, *Cd28*, *Tnfsf9*). These results are consistent with these m45 and m38 tumor resident T cells being highly activated/differentiated. Notably, cluster 7 which contains the highest numbers of m25 and m142 CD4 Tconv, had the most *Ifnγ* expression of any cluster, and showed high expression of co-stimulatory receptors *Cd28*, *Tnfrsf4* (OX40), *Tnfrsf18* (GITR), and *Cd40lg*, consistent with a strong effector phenotype. The top 5 differentially expressed genes in cluster 1 were *Havcr2* (TIM3), *Pdcd1* (PD1), *Rgs16*, *Lag3*, and *Cxcr6*, and in cluster 7 *Maf*, *Tnfrsf4*, *Glrx* and *Izumo1r* (Fig. S6D). Interestingly, glutaredoxin (*Glrx)* regulates redox homeostasis and cellular oxidative stress, which may help sustain T cell function in a stressful environment like the pancreatic TME.

### Immunization-induced T cells localize to tumors, but to a lesser extent than infection-induced T cells

Since MCMV-specific T cells preferentially localized to tumors in the absence iRGD, we sought to further investigate the underlying mechanism for this. Our first hypothesis was based on publications showing that CMV DNA is present, and the virus can reactivate/replicate, in glioblastoma tumors (37). In our experiments, mice are infected with MCMV and latency is established before tumor cell implantation. Therefore, if viral replication occurs directly in KPC tumor cells, reactivation from latency and subsequent infection of these cells would have to occur. Not unexpectedly, MCMV could productively infect KPC cells in culture, as shown using a MCMV-mCherry reporter virus (Fig 7A). Next, we tested whether we could detect MCMV DNA by qPCR in tumors implanted in infected mice. qPCR detected MCMV DNA in livers and spleens from day 4 acutely infected mice, as expected, but was undetectable in KPC1242 (Fig 7B) or KPC46 (Fig S7A) tumors.

**Figure 7.**
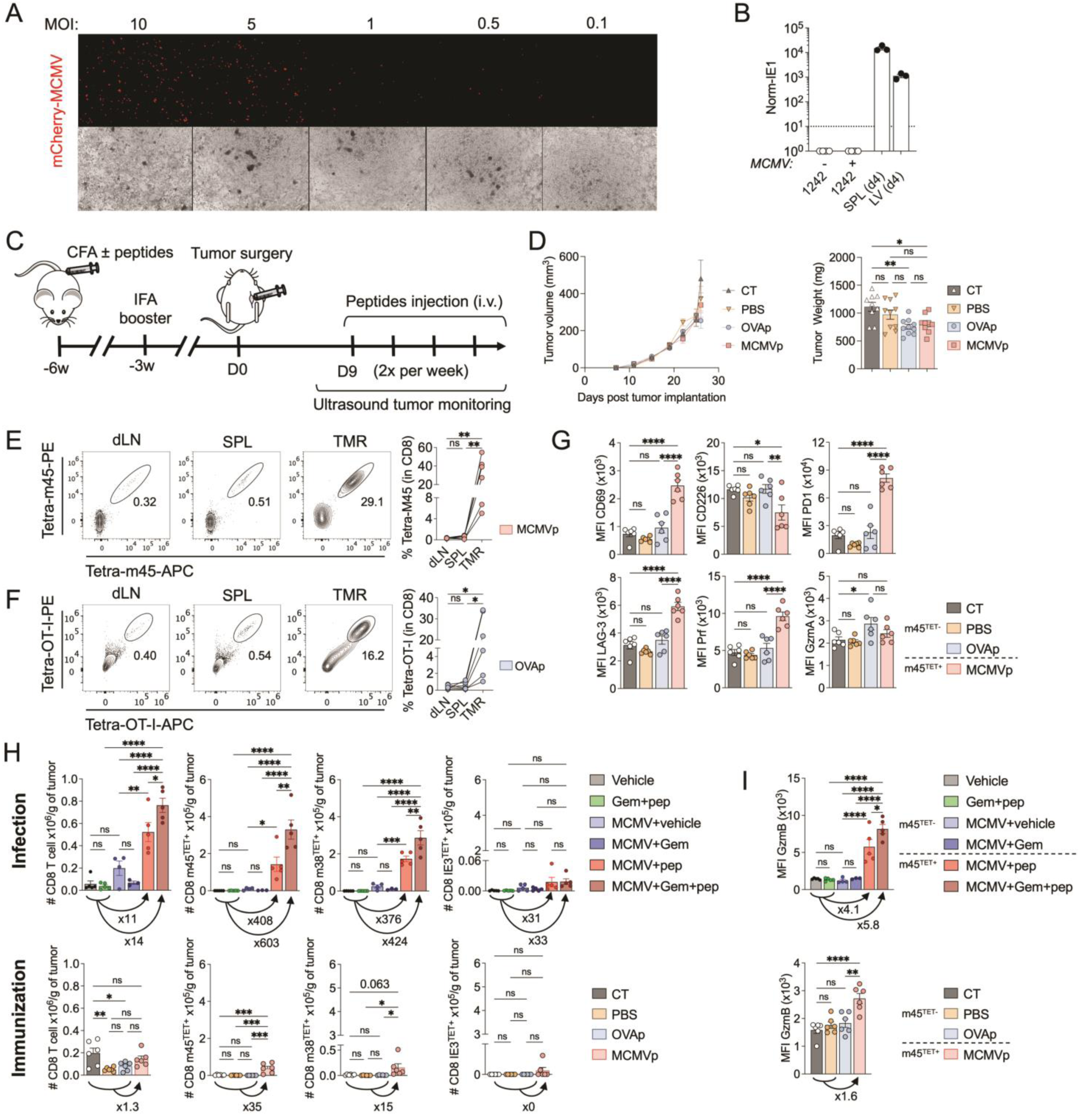
Immunization-induced T cells infiltrate tumors, but to a lesser extent than infection-induced T cells. **A**, KPC1242 infected *in vitro* with MCMV-mCherry for 48hr. **B**, IE1 DNA qPCR normalized to actin for KPC1242 tumor cells harvested from uninfected or MCMV infected animals after tumor implantation. Positive controls are spleen and liver from mice 4 dpi. **C**, Protocol for peptide immunization and tumor challenge. **D**, Tumor volume measured by ultrasound over time (left) and end point tumor weight (right) in the different treatment conditions after immunization (CT-Non immunized + MCMVp, PBS-Immunized with CFA/IFA no peptides, + MCMVp, OVAp-Immunized + OVA-I and OVA-IIp and MCMVp-Immunized + MCMVp). **E-I**, Mice from (D) were analyzed for T cell phenotype and function from the draining lymph node, spleen and tumor. **E**, Representative m45-tetramer binding among MCMVp-immunized mice, and proportions quantified over the 3 tissues. **F**, Representative OT-I-tetramer binding among OVAp-immunized mice, and proportions in the 3 tissues. **G**, Phenotype of tumor-infiltrating m45-specific CD8 T cells compared to m45-tetramer-negative. **H**, T cell quantification per gram of tumor, and fold differences seen between control and MCMVp treated groups after immunization and infection (top, from experiment shown in Fig.4). **I**, GzmB expression and fold increase comparing control and treated groups after immunization (bottom) and infection (top, from experiment shown in Fig.4). Results were compared by one-way ANOVA with Tukey correction. *P < 0.05; **P < 0.01; ***P < 0.001; ****P < 0.0001.

As these analyses showed that MCMV was not directly infecting the tumor and promoting virus-specific T cell recruitment, we next tested whether infection was even required for their tumor-residency. This was done by immunizing mice with MCMV or ovalbumin (OVA)-derived CD8 and CD4 T cell epitopes, implanting tumors and recalling these immunization-induced memory T cells with peptide injection once tumors were palpable (Fig. 7C). Tumor growth was monitored by ultrasound for 2 weeks and then mice were sacrificed for analysis of MCMV or OVA specific T cells from tumors, draining lymph nodes (dLN) and spleens. Contrary to MCMV infected mice, MCMVp therapy did not curtail tumor growth to nearly the same extent when memory T cells were induced by immunization. Recalling immunization-induced MCMV or OVA T cells had no impact on tumor volume and only a modest reduction of tumor weight was seen in both groups at the experimental end point (Fig. 7D). Somewhat unexpectedly, m45 and m38 CD8 T cells were found to be present in the tumors of the MCMVp immunized group (Fig. 7E), with fewer numbers of m25 and m142 CD4 Tconv (Fig. S7B).

These results proved that viral infection is not required for preferential accumulation of MCMV-specific T cells within the tumor in our experimental setup, and the same was seen for OVA-specific T cells (Fig. 7F). However, even though antigen-specific T cells infiltrated tumors after immunization, many fold fewer established tumor residence compared to those induced by infection: Total CD8 infiltration/gram of tumor increased 11-14 fold following infection compared to 1.3 fold after immunization, and m45 CD8 T cell number increased 408-603 fold in infected mice compared to ∼35 fold in immunized mice (Fig. 7H). This may explain the lack of tumor control observed in the context of immunization.

Phenotyping of immunization-induced tumor resident m45 CD8 T cells showed expression of CD69, PD1, LAG3, Prf and GzmB (Fig. 7G, I). Restimulation of CD45+ tumor cells with either OVA or MCMV CD8 T cell peptide epitopes also stimulated IFNγ production (Fig. S7C). However, the expression of activation and cytotoxic markers seen in these T cells was lower in immunized mice, as compared to infected mice after peptide recall of memory cells. Specifically, CD226, GzmA and GzmB weren’t increased to nearly the extent in immunization-induced m45 CD8 T cells after MCMVp therapy as seen in infected mice (e.g. GzmB expression by m45 CD8 T cells increased 1.6 fold compared to control group in the immunization protocol, compared to 4.1-5.8 fold in the infection protocol; Fig. 7I, S7D). Taken together, these results suggest that while viral infection is not required for preferential T cell tumor infiltration after MCMVp therapy, MCMV-specific memory T cells induced by infection infiltrate tumors in far greater numbers and seem to have higher cytotoxic potential, providing a rationale for the superior tumor growth control observed in the context of infection.

## Discussion

The 5-year survival for pancreatic cancer patients is 13% despite ongoing efforts to develop impactful new therapies. Pancreatic cancers are largely resistant to existing immunotherapies due to a highly immunosuppressive TME (8, 9) and a modest mutational burden (5–7), which results in a paucity of activated T cells that can reach and/or effectively kill cancer cells. In this work, we show that harnessing pre-existing antiviral memory T cells results in their massive tumor recruitment, substantial changes in the TME and ultimately reduced tumor growth and enhanced survival. We believe CMVp therapy represents a promising new approach to pancreatic cancer treatment in Human CMV (HCMV)-seropositive patients (∼80% of the world population). This therapy is (i) a relatively simple “off the shelf” T cell based therapy that is inexpensive compared to personalized medical approaches, (ii) shows delayed tumor growth and increased survival that we can enhance by combination with suboptimal doses of standard of care chemotherapy drugs, such as gemcitabine and (iii) there is pre-existing knowledge of the HCMV peptide sequences recognized by human CD4 and CD8T cells, some of which have been published by our laboratory. We have recently identified ∼200 new HCMV T cell epitopes (26), and are now testing the feasibility of using them for cancer treatment in humanized mouse models. These experiments should help identify HCMV epitopes that cover multiple haplotypes and are broadly applicable to CMVp therapy in people. Finally, (iv) the breadth and magnitude of CMV memory T cells is the largest known (>10% of circulating T cells), and harnessing this huge population via CMVp therapy may benefit from this fact.

While CMV-specific memory T cells are not the only ones “eligible” for this approach, our results suggest they may be a good choice. T cell expansion within the tumor following peptide immunization and subsequent recall was seen for both MCMV specific T cells and OVA specific OT-I CD8 T cells. However, the ability of these T cells to restrict tumor growth was not nearly as robust as that seen when MCMVp therapy was employed in latently infected mice. Perhaps this is due in part to the fact that peptide immunization does not induce the same “inflationary” MCMV-specific effector-memory CD8 T cell populations as seen during infection (38, 39). In addition, immunization-induced T cells infiltrated tumors at far lower numbers and expressed less GzmB when compared to those produced during infection (Fig. 7H, I and S7D).

scRNAseq analysis revealed substantial changes in the transcriptome of tumor epithelial cells after MCMVp therapy, leading to increased IFN-signaling pathways and decreased EMT, angiogenesis and Kras signaling. Finding that a technically simple peptide therapy can induce such dramatic changes by modifying the TME and increasing T cell-tumor cell interactions is encouraging, as it suggests multiple anti-tumor pathways are engaged. scRNAseq also allowed us to identify MCMV epitope-specific T cells based on their paired TCRαβ sequences. Notably, cluster 6 was increased after MCMVp therapy and contained highly activated CD8 T cells, but no MCMV-specific T cells. This suggests MCMVp therapy may unleash T cell responses specific for tumor neoantigens, thus broadening the scope of the antitumor response. Future sequencing of these TCRs should provide valuable information in this regard.

We initially co-injected the iRGD tumor-targeting peptide with MCMVp therapy, as we hypothesized it would specifically target the MCMV peptides to the tumor. Why did MCMVp therapy result in preferential tumor homing of MCMV specific T cells even in the absence of iRGD? We showed that MCMV DNA wasn’t present in tumor tissue, so TILs were not responding to infected cells. One possibility we have considered is that the enhanced macropinocytosis previously reported in pancreatic cancer cells (40–42) may supersede any requirements for iRGD in the uptake of systemically injected peptides. Finally, given the fact that multiple MCMV and OVA T cells induced by peptide immunization colonized the tumor, it is highly unlikely that molecular mimicry is the underlying mechanism for their trafficking to this site. Of note, while iRGD was not needed in this context, its usage as co-treatment with MCMVp therapy might still be necessary for the treatment of other tumor types, which remains to be tested.

In conclusion, this study demonstrates that systemic treatment with short T cell peptide epitopes can efficiently recall CMV memory T cells and result in their tumor localization and induction of an anti-tumor immune response. This strategy is a mutation agnostic approach that can be enhanced by combination therapy with low dose Gem and has the potential to benefit the majority of the population that harbor latent CMV. Importantly, our study reveals the benefit this approach provides in aggressive orthotopic pancreatic tumor models, where immunotherapy has failed, suggesting it may be reasonable for clinical translation.

## Materials and Methods

### Mice

Wild-type (WT) C57BL/6J (B6) mice and C57BL/6J x 129S1/SVlmJ (129) F1 hybrid (B6129SF1/J, Strain #:101043) were purchased from Jackson laboratories at 4-5 weeks of age, and infected or not with MCMV at 5-6 weeks of age. This study was carried out in strict accordance with the guidelines of Association for assessment and Accreditation of laboratory Animal Care (AAALAC) and National Institutes of Health (NIH). All animal protocols used in this study, were approved by the Institutional Animal Care and Use Committee (IACUC) of the University of California, San Diego and La Jolla Institute for immunology (LJI).

### MCMV infection

MCMV (Smith strain) was originally produced in 3T3 cells from cloned and sequenced BAC DNA (gift of B. Adler, (43)) and was then amplified *in vivo* in BALB/c mice, extracted from the salivary glands, and titrated on MEF (murine embryonic fibroblast) cells (17)..5-6-weeks old mice were infected i.p. with 1 x 10^4^ pfu in 100µL of PBS. The mCherry cytomegalovirus was kindly provided by Dr. Lars Dolken.

### Pancreatic tumor models

MCMV infected mice (2-3 months post infection) and age matched uninfected controls were orthotopically injected with KPC1242 (syngeneic with B6 mice) or KPC46 (syngeneic with B6129SF1/J) pancreatic cancer cells. These cells were cultured *in vitro* in RPMI with 10% Fetal Bovine Serum (FBS) and 1% penicillin/streptomycin. Once confluent, they were trypsinized, washed and resuspended in Matrigel (Corning Matrigel Matrix) for injection (5,000 KPC1242 or KPC46 tumor cells in 20ul of matrigel, per mouse). Tumor growth was monitored initially by palpation and after day 8 post injection, mice were subjected to ultrasound (Convex L20 HD3, Clarius) twice a week, (under isofluorane) for tumor growth monitoring. Tumor volume was calculated using the ellipsoid volume calculation formula. In addition, mouse weights were taken twice a week together with health checks. Mice were enrolled in treatment studies and randomly assigned to the different treatment conditions once most tumors were visible by ultrasound, usually around 9-11 d.p.i. For survival studies, the euthanasia criteria included mice loosing 20% or more of their initial weight or having tumors reaching 2000mm^3^ or more plus any evident signs of distress.

### Treatment Studies

MCMV peptides (CD4T: m09_133-147_; m25_409-423_; m142_24-38_; CD8T: IE3_416-423_; m38_316-323_; m45_985-993_) were purchased from Mimotopes (Australia), and ovalbumin peptides (OVA_257-264_; OVA_323-339_) were from Eurogentec-Anaspec, CA, USA. iRGD (Certepetide) was acquired from LISATA Therapeutics. Mice were treated twice a week by retro-orbital (RO) injections of the indicated peptides: Dose A (50µg CD4p/1µg CD8p); Dose B (50µg CD4p/0.1µg CD8p); Dose C (10µg CD4p/3µg CD8p); Dose D (10µg CD4p/1µg CD8p). When iRGD was utilized, mice were first injected with iRGD (300µg) in one eye followed by the mix of MCMV peptides 5min later in the other eye. When indicated, mice were intraperitoneally injected with anti-PD1 (10mg/kg twice a week, Bioxcell) or anti-IL10R (200μg/ul, once a week, Bioxcell) antibodies, or gemcitabine (5mg/kg, twice a week, Selleck Chem). Control mice were injected RO with vehicle (PBS/DMSO equivalent to the amount in the MCMV peptide mix) and saline or rat IgG controls for αPD1 and αIL10R blocking antibodies IP.

### Mice Immunization

8-week old mice were immunized subcutaneously at the base of the tail with 200µL of an emulsion containing 50µg of each indicated peptide (MCMVp or OVAp) and CFA (0.5mg/mL final; Sigma). Booster immunizations in containing peptides + IFA were performed 3 weeks later. Mice were tumor challenged 3 weeks later as previously described.

### Histology/Immunohistology

Upon harvesting, tumors were fixed in 10% Formalin for 24-48 hours and kept in 70% ethanol until paraffin embedded. Immunohistochemistry was performed as previously described (18). In short, FFPE sections were de-paraffinized, and subjected to steam heat mediated antigen retrieval using low pH buffer (eBioscience). Blocking was performed with 5% donkey and 5% goat serum. The sections were analyzed for cleaved Caspase 3 (1:250) (Cat.no. 9579S, Cell Signaling), and CD3 (1:250) (Cat.no. 11089, Abcam) and kept at 4 °C overnight and incubated with their respective secondary antibodies. Histological and immunofluorescence imaging was performed using a ZEISS AxioScan Z1 automated slide scanning microscope equipped with a 20x objective (NA 0.8). For brightfield imaging of H&E-stained specimens, an LED illumination system was employed in conjunction with a Hitachi HV-FL202SCL camera. Immunofluorescence imaging was conducted using a Colibri7 LED illumination system and ZEISS single-band filter sets (filter sets 49, 38, 43, and 50). Fluorescence images were captured using a Hamamatsu Orca Flash 4.0 v2 camera operating in 16-bit mode (Microscopy core, LJI). Staining was quantified using the Qpath software. Hematoxilyn and Eosin staining was performed also as previously described (18) and necrosis scores were assessed by a histopathologist blinded to the different treatment conditions.

### FACS analysis

Tumors were minced into small pieces and incubated at 37°C with rotation for 30 minutes in 5 mL of digestion buffer consisting of DMEM high glucose, 10% Gentle Collagenase/Hyaluronidase (GCH, STEM CELL), 10% FBS, and 10% DNaseI (1 mg/mL stock, Roche). Samples were then added on top of a 70-μm filter and smashed with the back of a syringe while adding 5 mL of PBS 2% FBS. Flow through was spun at 300*g* for 10 minutes and subjected to RBC lysis (Pharmalyse, BD, Biosciences). Cells were counted and subjected to magnetic separation of tumor infiltrating lymphocytes using Easysep TIL kit (STEM CELL). Spleen and liver cells were harvested and processed as previously described (44) and analyzed on an LSR-II Fortessa X20 (BD) or a Cytek Aurora. Dead cells were excluded by staining with LD blue (Invitrogen). Surface staining was performed by diluting antibodies in Brilliant Stain buffer (Invitrogen). Fixation/permeabilization was done with Foxp3/Transcription Factor Staining Buffer Set (eBioscience). For antibodies used see supplementary materials.

For tetramer-staining, NIH-provided biotinylated monomers were tetramerized in the lab with streptavidin-APC or -PE, and subsequently used to stain the cells as previously described (43). Briefly, tetramer staining was done at RT for 1.5-2h. Live/dead and surface/intracellular staining were done subsequently. Data was analyzed using the FlowJo software.

### Cytokine stimulation

Total liver or purified CD45+ cells from the tumors were incubated for 4h at 37°C 5% C02 in 96-well plates in the presence of the indicated peptides (5µg/mL for each peptide) and GolgiPlug (Brefeldin A; eBioscience). Control cells were incubated in presence of GolgiPlug alone. Cells were then stained as described above.

### Single-cell RNA-seq

Total live-tumor cells (n=3 per group) or total tumor-infiltrating T cells (n=3 per group) (some samples were a combination of 2 tumors) were sorted from the indicated group, counted, and processed for subsequent 5’scRNAseq using Chromium GEM-X single cell 5’ v3 gene expression kit. See supplementary material for analysis pipeline.

### Data analysis

Data is presented as means ± SEM. The GraphPad Prism statistical package was used for statistical analyses (Graph-Pad Software, Inc.). Results were compared by one-way ANOVA with Tukey correction, and repeated measures by two-way A NOVA with Sidak correction. Results were considered statistically significant when P < 0.05 and are indicated in the figures as follows: *P < 0.05; **P < 0.01; ***P < 0.001; ****P < 0.0001.

## Acknowledgments

NGS, Flow Cytometry, Microscopy, and Histological cores facilities at the La Jolla Institute. Microscopy and flow cytometry cores at Moores Cancer Center.

NIH for providing MHC-I/II biotinylated monomer. This work was supported by Foundation for a Better World grant 001 to T.H.M. and NCI grant 1R21CA286198 to T.H.M. and C.A.B. The funders had no role in study design, data collection and analysis, decision to publish, or preparation of the manuscript.

## Data availability

The data and materials are available on requests.

## Abbreviations

Gem: gemcitabine
HCMV: human cytomegalovirus
KPC: kras-pdx1-cre pancreatic adenocarcinoma
MCMV: murine cytomegalovirus
MCMVp: MCMV-peptide
Tconv: conventional CD4 T cell (Foxp3^−^)
TILs: tumor infiltrating lymphocytes
TME: tumor microenvironment
Treg: regulatory CD4 T cell (Foxp3^+^)

## References

1. Schadendorf, D., F. S. Hodi, C. Robert, J. S. Weber, K. Margolin, O. Hamid, D. Patt, T.-T. Chen, D. M. Berman, and J. D. Wolchok. 2015. Pooled Analysis of Long-Term Survival Data From Phase II and Phase III Trials of Ipilimumab in Unresectable or Metastatic Melanoma. JCO 33: 1889–1894.

2. Reck, M., D. Rodríguez-Abreu, A. G. Robinson, R. Hui, T. Csőszi, A. Fülöp, M. Gottfried, N. Peled, A. Tafreshi, S. Cuffe, M. O’Brien, S. Rao, K. Hotta, M. A. Leiby, G. M. Lubiniecki, Y. Shentu, R. Rangwala, and J. R. Brahmer. 2016. Pembrolizumab versus Chemotherapy for PD-L1–Positive Non–Small-Cell Lung Cancer. N Engl J Med 375: 1823–1833.

3. Garon, E. B., N. A. Rizvi, R. Hui, N. Leighl, A. S. Balmanoukian, J. P. Eder, A. Patnaik, C. Aggarwal, M. Gubens, L. Horn, E. Carcereny, M.-J. Ahn, E. Felip, J.-S. Lee, M. D. Hellmann, O. Hamid, J. W. Goldman, J.-C. Soria, M. Dolled-Filhart, R. Z. Rutledge, J. Zhang, J. K. Lunceford, R. Rangwala, G. M. Lubiniecki, C. Roach, K. Emancipator, and L. Gandhi. 2015. Pembrolizumab for the Treatment of Non–Small-Cell Lung Cancer. N Engl J Med 372: 2018–2028.

4. Wei, S. C., C. R. Duffy, and J. P. Allison. 2018. Fundamental Mechanisms of Immune Checkpoint Blockade Therapy. Cancer Discovery 8: 1069–1086.

5. Australian Pancreatic Cancer Genome Initiative, ICGC Breast Cancer Consortium, ICGC MMML-Seq Consortium, ICGC PedBrain, L. B. Alexandrov, S. Nik-Zainal, D. C. Wedge, S. A. J. R. Aparicio, S. Behjati, A. V. Biankin, G. R. Bignell, N. Bolli, A. Borg, A.-L. Børresen-Dale, S. Boyault, B. Burkhardt, A. P. Butler, C. Caldas, H. R. Davies, C. Desmedt, R. Eils, J. E. Eyfjörd, J. A. Foekens, M. Greaves, F. Hosoda, B. Hutter, T. Ilicic, S. Imbeaud, M. Imielinski, N. Jäger, D. T. W. Jones, D. Jones, S. Knappskog, M. Kool, S. R. Lakhani, C. López-Otín, S. Martin, N. C. Munshi, H. Nakamura, P. A. Northcott, M. Pajic, E. Papaemmanuil, A. Paradiso, J. V. Pearson, X. S. Puente, K. Raine, M. Ramakrishna, A. L. Richardson, J. Richter, P. Rosenstiel, M. Schlesner, T. N. Schumacher, P. N. Span, J. W. Teague, Y. Totoki, A. N. J. Tutt, R. Valdés-Mas, M. M. van Buuren, L. van’t Veer, A. Vincent-Salomon, N. Waddell, L. R. Yates, J. Zucman-Rossi, P. Andrew Futreal, U. McDermott, P. Lichter, M. Meyerson, S. M. Grimmond, R. Siebert, E. Campo, T. Shibata, S. M. Pfister, P. J. Campbell, and M. R. Stratton. 2013. Signatures of mutational processes in human cancer. Nature 500: 415–421.

6. Schumacher, T. N., and R. D. Schreiber. 2015. Neoantigens in cancer immunotherapy. Science 348: 69–74.

7. Evans, R. A., M. S. Diamond, A. J. Rech, T. Chao, M. W. Richardson, J. H. Lin, D. L. Bajor, K. T. Byrne, B. Z. Stanger, J. L. Riley, N. Markosyan, R. Winograd, and R. H. Vonderheide. 2016. Lack of immunoediting in murine pancreatic cancer reversed with neoantigen. JCI Insight 1.

8. Hiraoka, N., K. Onozato, T. Kosuge, and S. Hirohashi. 2006. Prevalence of FOXP3+ Regulatory T Cells Increases During the Progression of Pancreatic Ductal Adenocarcinoma and Its Premalignant Lesions. Clinical Cancer Research 12: 5423–5434.

9. Ino, Y., R. Yamazaki-Itoh, K. Shimada, M. Iwasaki, T. Kosuge, Y. Kanai, and N. Hiraoka. 2013. Immune cell infiltration as an indicator of the immune microenvironment of pancreatic cancer. Br J Cancer 108: 914–923.

10. Brahmer, J. R., S. S. Tykodi, L. Q. M. Chow, W.-J. Hwu, S. L. Topalian, P. Hwu, C. G. Drake, L. H. Camacho, J. Kauh, K. Odunsi, H. C. Pitot, O. Hamid, S. Bhatia, R. Martins, K. Eaton, S. Chen, T. M. Salay, S. Alaparthy, J. F. Grosso, A. J. Korman, S. M. Parker, S. Agrawal, S. M. Goldberg, D. M. Pardoll, A. Gupta, and J. M. Wigginton. 2012. Safety and Activity of Anti–PD-L1 Antibody in Patients with Advanced Cancer. N Engl J Med 366: 2455–2465.

11. Foley, K., V. Kim, E. Jaffee, and L. Zheng. 2016. Current progress in immunotherapy for pancreatic cancer. Cancer Letters 381: 244–251.

12. Royal, R. E., C. Levy, K. Turner, A. Mathur, M. Hughes, U. S. Kammula, R. M. Sherry, S. L. Topalian, J. C. Yang, I. Lowy, and S. A. Rosenberg. 2010. Phase 2 Trial of Single Agent Ipilimumab (Anti-CTLA-4) for Locally Advanced or Metastatic Pancreatic Adenocarcinoma. Journal of Immunotherapy 33: 828–833.

13. Balachandran, V. P., M. Łuksza, J. N. Zhao, V. Makarov, J. A. Moral, R. Remark, B. Herbst, G. Askan, U. Bhanot, Y. Senbabaoglu, D. K. Wells, C. I. O. Cary, O. Grbovic-Huezo, M. Attiyeh, B. Medina, J. Zhang, J. Loo, J. Saglimbeni, M. Abu-Akeel, R. Zappasodi, N. Riaz, M. Smoragiewicz, Z. L. Kelley, O. Basturk, Australian Pancreatic Cancer Genome Initiative, Garvan Institute of Medical Research, Prince of Wales Hospital, Royal North Shore Hospital, University of Glasgow, St Vincent’s Hospital, QIMR Berghofer Medical Research Institute, University of Melbourne, Centre for Cancer Research, University of Queensland, Institute for Molecular Bioscience, Bankstown Hospital, Liverpool Hospital, Royal Prince Alfred Hospital, Chris O’Brien Lifehouse, Westmead Hospital, Fremantle Hospital, St John of God Healthcare, Royal Adelaide Hospital, Flinders Medical Centre, Envoi Pathology, Princess Alexandria Hospital, Austin Hospital, Johns Hopkins Medical Institutes, ARC-Net Centre for Applied Research on Cancer, M. Gönen, A. J. Levine, P. J. Allen, D. T. Fearon, M. Merad, S. Gnjatic, C. A. Iacobuzio-Donahue, J. D. Wolchok, R. P. DeMatteo, T. A. Chan, B. D. Greenbaum, T. Merghoub, and S. D. Leach. 2017. Identification of unique neoantigen qualities in long-term survivors of pancreatic cancer. Nature 551: 512–516.

14. Rosato, P. C., S. Wijeyesinghe, J. M. Stolley, C. E. Nelson, R. L. Davis, L. S. Manlove, C. A. Pennell, B. R. Blazar, C. C. Chen, M. A. Geller, V. Vezys, and D. Masopust. 2019. Virus-specific memory T cells populate tumors and can be repurposed for tumor immunotherapy. Nat Commun 10: 567.

15. Çuburu, N., L. Bialkowski, S. M. Pontejo, S. K. Sethi, A. T. F. Bell, R. Kim, C. D. Thompson, D. R. Lowy, and J. T. Schiller. 2022. Harnessing anti-cytomegalovirus immunity for local immunotherapy against solid tumors. Proc. Natl. Acad. Sci. U.S.A. 119: e2116738119.

16. Teesalu, T., K. N. Sugahara, V. R. Kotamraju, and E. Ruoslahti. 2009. C-end rule peptides mediate neuropilin-1-dependent cell, vascular, and tissue penetration. Proc. Natl. Acad. Sci. U.S.A. 106: 16157–16162.

17. Sugahara, K. N., T. Teesalu, P. P. Karmali, V. R. Kotamraju, L. Agemy, D. R. Greenwald, and E. Ruoslahti. 2010. Coadministration of a Tumor-Penetrating Peptide Enhances the Efficacy of Cancer Drugs. Science 328: 1031–1035.

18. Hurtado de Mendoza, T., E. S. Mose, G. P. Botta, G. B. Braun, V. R. Kotamraju, R. P. French, K. Suzuki, N. Miyamura, T. Teesalu, E. Ruoslahti, A. M. Lowy, and K. N. Sugahara. 2021. Tumor-penetrating therapy for β5 integrin-rich pancreas cancer. Nature Communications 12: 1541.

19. Sugahara, K. N., G. B. Braun, T. H. de Mendoza, V. R. Kotamraju, R. P. French, A. M. Lowy, T. Teesalu, and E. Ruoslahti. 2015. Tumor-Penetrating iRGD Peptide Inhibits Metastasis. Molecular Cancer Therapeutics 14: 120–128.

20. Liu, X., P. Lin, I. Perrett, J. Lin, Y.-P. Liao, C. H. Chang, J. Jiang, N. Wu, T. Donahue, Z. Wainberg, A. E. Nel, and H. Meng. 2017. Tumor-penetrating peptide enhances transcytosis of silicasome-based chemotherapy for pancreatic cancer. Journal of Clinical Investigation 127: 2007–2018.

21. Song, W., M. Li, Z. Tang, Q. Li, Y. Yang, H. Liu, T. Duan, H. Hong, and X. Chen. 2012. Methoxypoly(ethylene glycol) *-block-* Poly(L-glutamic acid)-Loaded Cisplatin and a Combination With iRGD for the Treatment of Non-Small-Cell Lung Cancers. Macromol. Biosci. 12: 1514–1523.

22. Deng, C., M. Jia, G. Wei, T. Tan, Y. Fu, H. Gao, X. Sun, Q. Zhang, T. Gong, and Z. Zhang. 2017. Inducing Optimal Antitumor Immune Response through Coadministering iRGD with Pirarubicin Loaded Nanostructured Lipid Carriers for Breast Cancer Therapy. Mol. Pharmaceutics 14: 296–309.

23. Sha, H., R. Li, X. Bian, Q. Liu, C. Xie, X. Xin, W. Kong, X. Qian, X. Jiang, W. Hu, and B. Liu. 2015. A tumor-penetrating recombinant protein anti-EGFR-iRGD enhance efficacy of paclitaxel in 3D multicellular spheroids and gastric cancer in vivo. European Journal of Pharmaceutical Sciences 77: 60–72.

24. Ding, N., Z. Zou, H. Sha, S. Su, H. Qian, F. Meng, F. Chen, S. Du, S. Zhou, H. Chen, L. Zhang, J. Yang, J. Wei, and B. Liu. 2019. iRGD synergizes with PD-1 knockout immunotherapy by enhancing lymphocyte infiltration in gastric cancer. Nat Commun 10: 1336.

25. Agemy, L., D. Friedmann-Morvinski, V. R. Kotamraju, L. Roth, K. N. Sugahara, O. M. Girard, R. F. Mattrey, I. M. Verma, and E. Ruoslahti. 2011. Targeted nanoparticle enhanced proapoptotic peptide as potential therapy for glioblastoma. Proc. Natl. Acad. Sci. U.S.A. 108: 17450–17455.

26. Dhanwani, R., S. K. Dhanda, J. Pham, G. P. Williams, J. Sidney, A. Grifoni, G. Picarda, C. S. Lindestam Arlehamn, A. Sette, and C. A. Benedict. 2021. Profiling Human Cytomegalovirus-Specific T Cell Responses Reveals Novel Immunogenic Open Reading Frames. J Virol 95: e0094021.

27. Picarda, G., and C. A. Benedict. 2018. Cytomegalovirus: Shape-Shifting the Immune System. J Immunol 200: 3881–3889.

28. Suzuki, E., V. Kapoor, A. S. Jassar, L. R. Kaiser, and S. M. Albelda. 2005. Gemcitabine selectively eliminates splenic Gr-1+/CD11b+ myeloid suppressor cells in tumor-bearing animals and enhances antitumor immune activity. Clin Cancer Res 11: 6713–6721.

29. Sasso, M. S., G. Lollo, M. Pitorre, S. Solito, L. Pinton, S. Valpione, G. Bastiat, S. Mandruzzato, V. Bronte, I. Marigo, and J.-P. Benoit. 2016. Low dose gemcitabine-loaded lipid nanocapsules target monocytic myeloid-derived suppressor cells and potentiate cancer immunotherapy. Biomaterials 96: 47–62.

30. Eriksson, E., J. Wenthe, S. Irenaeus, A. Loskog, and G. Ullenhag. 2016. Gemcitabine reduces MDSCs, tregs and TGFβ-1 while restoring the teff/treg ratio in patients with pancreatic cancer. J Transl Med 14: 282.

31. Kissick, H., and R. Ahmed. 2022. New epigenetic regulators of T cell exhaustion. Cancer Cell 40: 708–710.

32. Kalluri, R. 2016. The biology and function of fibroblasts in cancer. Nat Rev Cancer 16: 582–598.

33. Du, Y., Y. Lin, L. Gan, S. Wang, S. Chen, C. Li, S. Hou, B. Hu, B. Wang, Y. Ye, and Z. Shen. 2024. Potential crosstalk between SPP1 + TAMs and CD8 + exhausted T cells promotes an immunosuppressive environment in gastric metastatic cancer. J Transl Med 22: 158.

34. Wu, J., Y. Shen, G. Zeng, Y. Liang, and G. Liao. 2024. SPP1+ TAM subpopulations in tumor microenvironment promote intravasation and metastasis of head and neck squamous cell carcinoma. Cancer Gene Ther 31: 311–321.

35. Smith, C. J., V. Venturi, M. F. Quigley, H. Turula, E. Gostick, K. Ladell, B. J. Hill, D. Himelfarb, K. M. Quinn, H. Y. Greenaway, T. H. Y. Dang, R. A. Seder, D. C. Douek, A. B. Hill, M. P. Davenport, D. A. Price, and C. M. Snyder. 2020. Stochastic Expansions Maintain the Clonal Stability of CD8+ T Cell Populations Undergoing Memory Inflation Driven by Murine Cytomegalovirus. J Immunol 204: 112–121.

36. Dash, P., A. J. Fiore-Gartland, T. Hertz, G. C. Wang, S. Sharma, A. Souquette, J. C. Crawford, E. B. Clemens, T. H. O. Nguyen, K. Kedzierska, N. L. La Gruta, P. Bradley, and P. G. Thomas. 2017. Quantifiable predictive features define epitope-specific T cell receptor repertoires. Nature 547: 89–93.

37. Krenzlin, H., P. Behera, V. Lorenz, C. Passaro, M. Zdioruk, M. O. Nowicki, K. Grauwet, H. Zhang, M. Skubal, H. Ito, R. Zane, M. Gutknecht, M. B. Griessl, F. Ricklefs, L. Ding, S. Peled, A. Rooj, C. D. James, C. S. Cobbs, C. H. Cook, E. A. Chiocca, and S. E. Lawler. 2019. Cytomegalovirus promotes murine glioblastoma growth via pericyte recruitment and angiogenesis. Journal of Clinical Investigation 129: 1671–1683.

38. Munks, M. W., K. S. Cho, A. K. Pinto, S. Sierro, P. Klenerman, and A. B. Hill. 2006. Four Distinct Patterns of Memory CD8 T Cell Responses to Chronic Murine Cytomegalovirus Infection. The Journal of Immunology 177: 450–458.

39. Panagioti, E., A. Redeker, S. Van Duikeren, K. L. Franken, J. W. Drijfhout, S. H. Van Der Burg, and R. Arens. 2016. The Breadth of Synthetic Long Peptide Vaccine-Induced CD8+ T Cell Responses Determines the Efficacy against Mouse Cytomegalovirus Infection. PLoS Pathog 12: e1005895.

40. Commisso, C., S. M. Davidson, R. G. Soydaner-Azeloglu, S. J. Parker, J. J. Kamphorst, S. Hackett, E. Grabocka, M. Nofal, J. A. Drebin, C. B. Thompson, J. D. Rabinowitz, C. M. Metallo, M. G. Vander Heiden, and D. Bar-Sagi. 2013. Macropinocytosis of protein is an amino acid supply route in Ras-transformed cells. Nature 497: 633–637.

41. Qiu, Z., W. Liu, Q. Zhu, K. Ke, Q. Zhu, W. Jin, S. Yu, Z. Yang, L. Li, X. Sun, S. Ren, Y. Liu, Z. Zhu, J. Zeng, X. Huang, Y. Huang, L. Wei, M. Ma, J. Lu, X. Chen, Y. Mou, T. Xie, and X. Sui. 2022. The Role and Therapeutic Potential of Macropinocytosis in Cancer. Front. Pharmacol. 13: 919819.

42. Andreasson, C., D. Ansari, F. Ekbom, and R. Andersson. 2021. Macropinocytosis: the Achilles’ heel of pancreatic cancer? Scandinavian Journal of Gastroenterology 56: 177–179.

43. Brunel, S., G. Picarda, A. Gupta, R. Ghosh, B. McDonald, R. El Morabiti, W. Jiang, J. A. Greenbaum, B. Adler, G. Seumois, M. Croft, P. Vijayanand, and C. A. Benedict. 2024. Late-rising CD4 T cells resolve mouse cytomegalovirus persistent replication in the salivary gland. PLoS Pathog 20: e1011852.

44. Picarda, G., R. Ghosh, B. McDonald, S. Verma, N. Thiault, R. El Morabiti, T. S. Griffith, and C. A. Benedict. 2019. Cytomegalovirus Evades TRAIL-Mediated Innate Lymphoid Cell 1 Defenses. J Virol 93: e00617–19.

