## Supplementary Figures and Methods for "Redirecting cytomegalovirus immunity against pancreas cancer for immunotherapy"

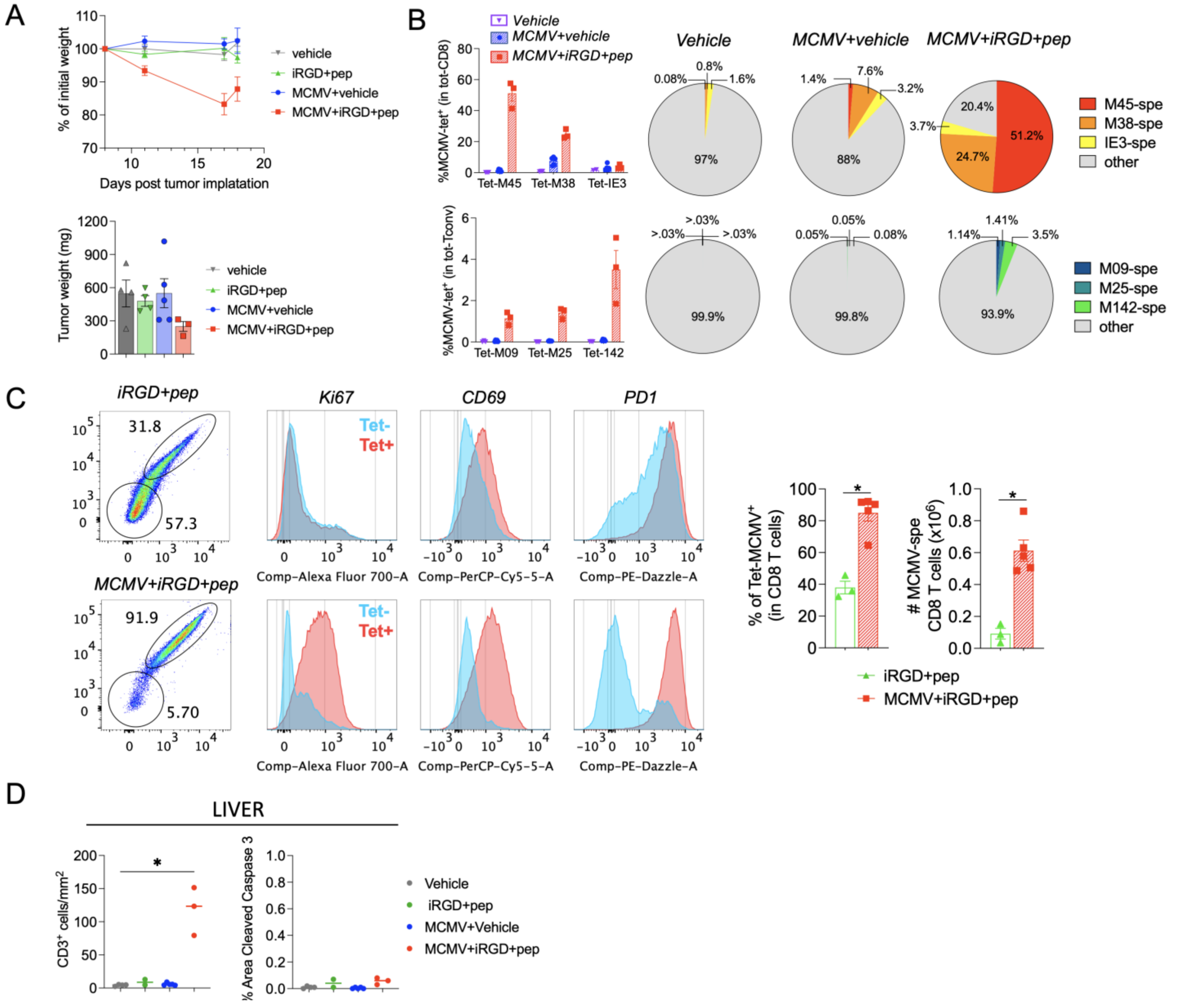


**Suppl Figure 1. MCMVp treatment induces a marked expansion of virus specific T cells. A**, Normalized weights from mice shown in Fig. 1B over the course of treatment with 50 µg of each MCMV peptide epitope (top). Tumor weights in the different treatment groups at end point (18 days post tumor implantation) (bottom). **B,** MCMV-tetramer quantification in the spleen of the mice shown in Fig. 1B. **C,** MCMV-tetramer quantification and phenotype of CD8 T cells in tumors of infected or uninfected mice treated with iRGD+MCMVp. **D,** Graphs showing the quantification of immunofluorescent staining of liver sections with CD3 and cleaved caspase 3 to measure T cell infiltration and apoptosis. Results were compared by Mann-Whitney analysis. *P < 0.05; **P < 0.01; ***P < 0.001; ****P < 0.0001.

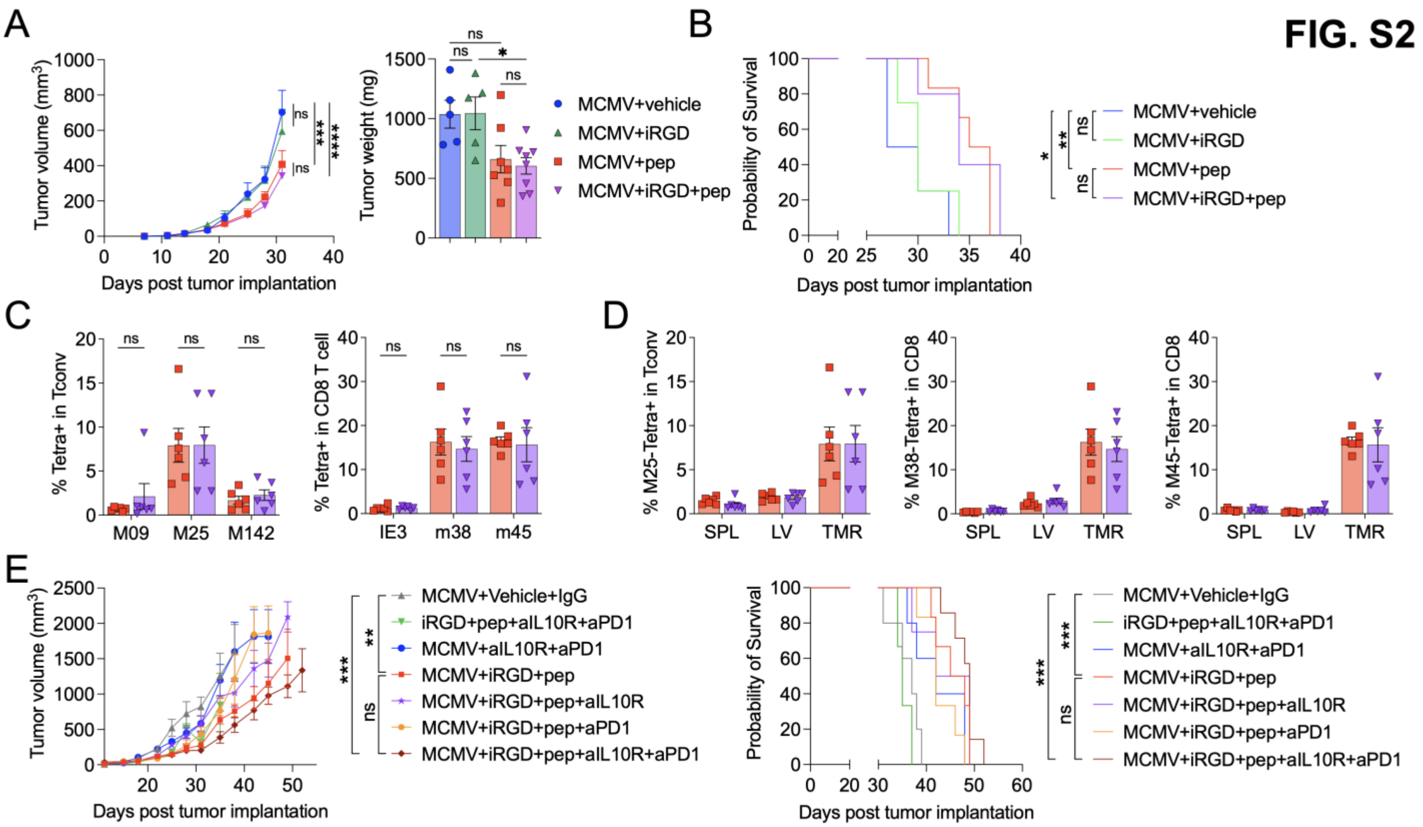


**Suppl Figure 2. MCMV-specific T cells preferentially localize within tumors in the absence of iRGD. A**, Growth curve of KPC1242 tumors over time (left) and tumor weight at endpoint (right). Dose B was used in combination or not with iRGD. **B**, Kaplan-Meier survival graph with a similar protocol as shown in (Fig.2A) using Dose A. **C-D**, MCMV specific T cell analysis from the spleen, liver and tumor at endpoint from the mice shown in (A). **C**, Percentage of each tetramer among total Tconv (left) or CD8 T cells (right) in the tumor. **D**, Percentage of the 3 dominant MCMV-specific T cells (m25, m38, and m45) in the 3 tissues tested. **E**, Assessment of combination therapy to boost MCMVp treatment with anti-IL10R and/or anti-PD1. Tumor growth over time (left) and Kaplan-Meier survival graph (right). Results were compared by one-way ANOVA with Tukey correction, and repeated measures by two-way A NOVA with Sidak correction. $^*P < 0.05$; $^{***}P < 0.001$; $^{****}P < 0.0001$.

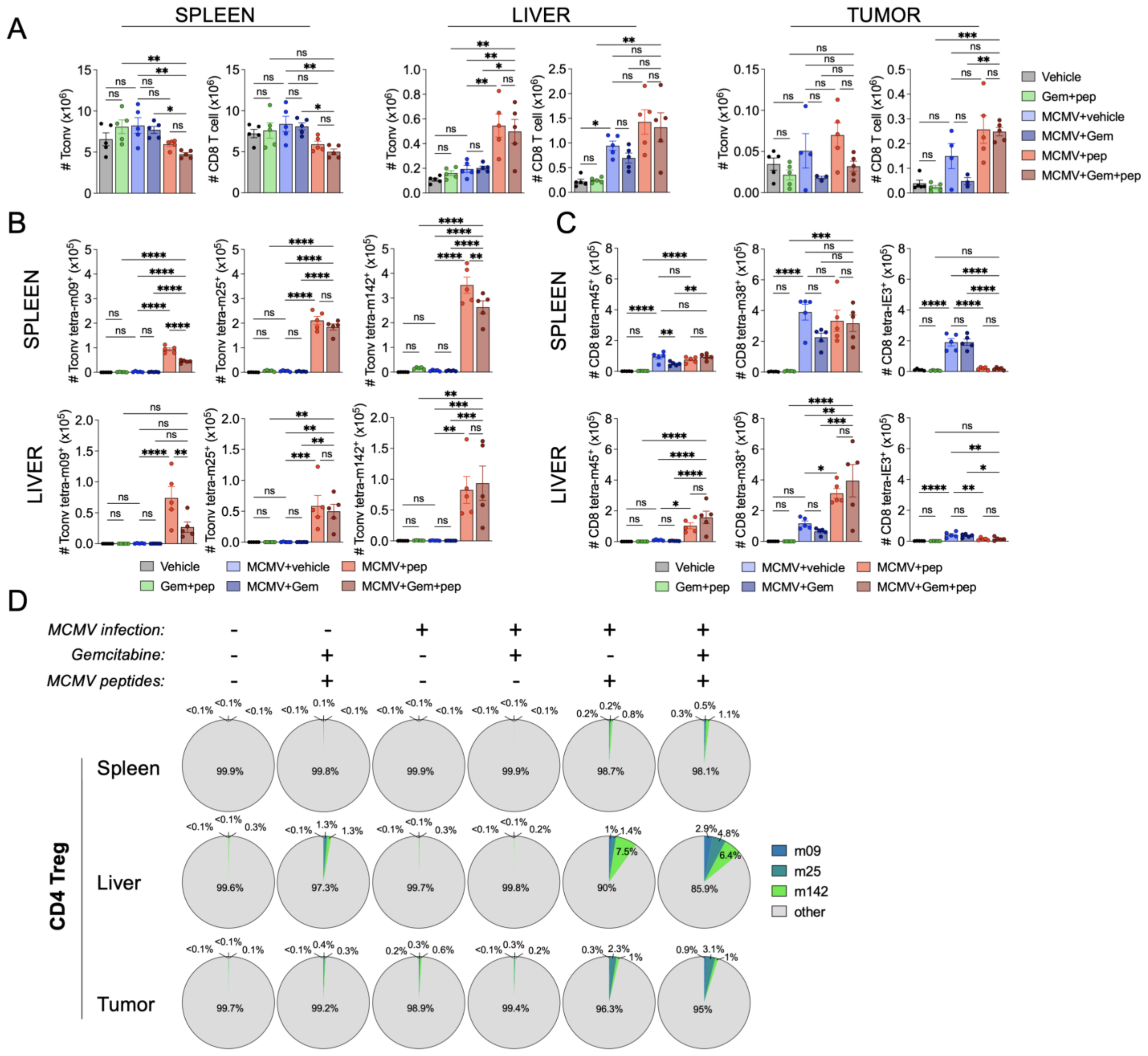


**Suppl Figure 3. MCMV-specific T cells expand following treatment, and preferentially localize to the tumor.** Mice were tumor challenged and treated as described in Figure 2B, to analyze spleen, liver, and tumor T cell infiltrates. (n=5/group). **A**, Absolute number of CD4 Tconv and CD8 T cell quantification at endpoint in the 3 tissues analyzed. **B-C**, Absolute number of MCMV-specific T cells at endpoint in the spleen and liver for Tconv (B) and CD8 T cells (C). **D**, Proportion of MCMV-specific FoxP3+ Tregs among total Tregs in the 3 tissues analyzed. Results were compared by one-way ANOVA with Tukey correction. *$P < 0.05$; **$P < 0.01$; ***$P < 0.001$; ****$P < 0.0001$.

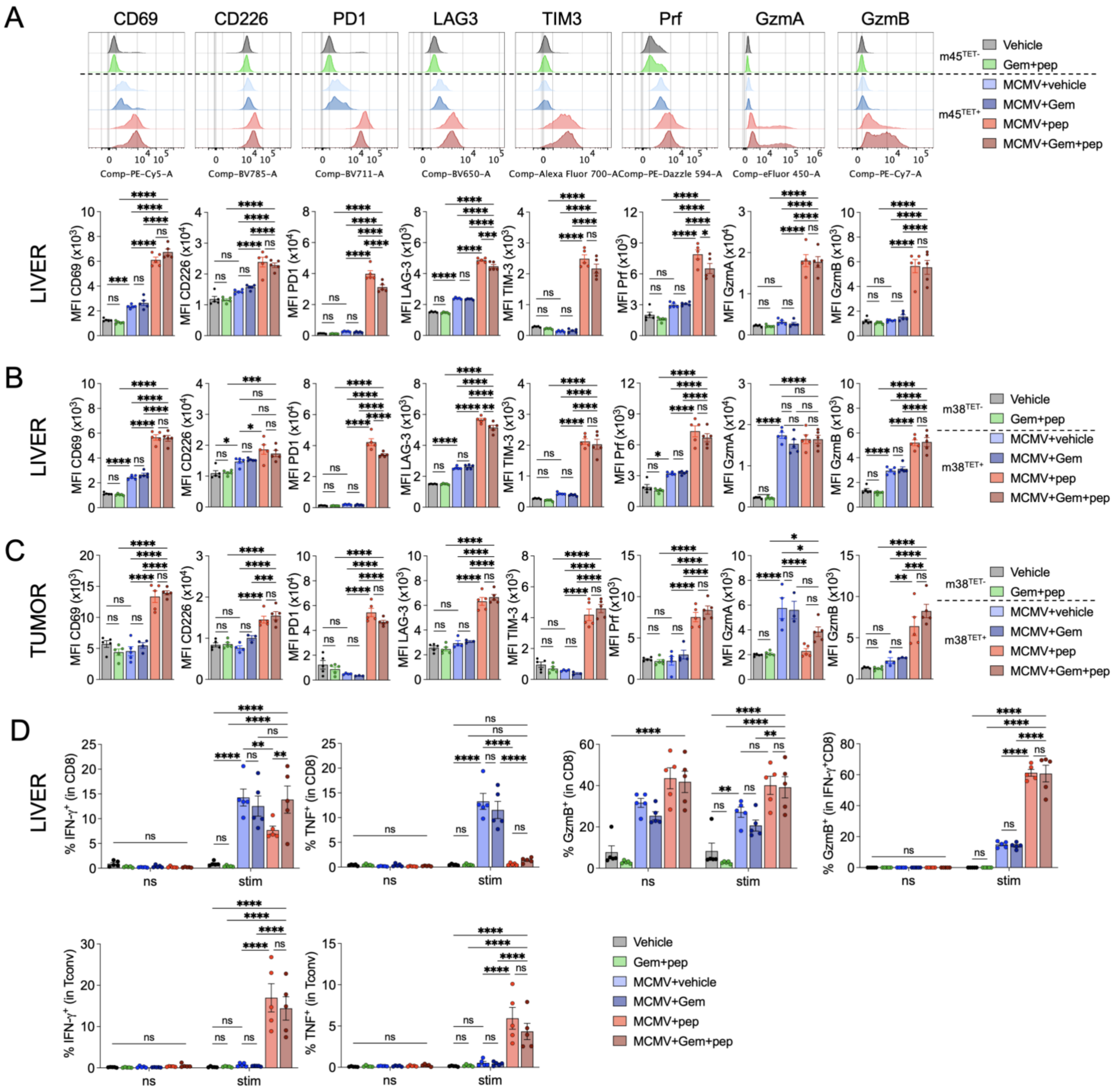


**Suppl Figure 4. MCMV-specific T cells are highly activated, cytotoxic and cytokine producers.** Immunophenotyping and cytokine production analysis of the TILs described in Fig 3. (n=5/group). **A**, Representative histograms of the m45-specific CD8 T cells in the liver stained with the indicated markers (top) and mean fluorescence intensity (MFI) quantification (bottom). **B-C**, Same analysis for m38-specific CD8 T cells in the liver (B) and tumor (C). **D**, Total liver cells were incubated with the 6 MCMV peptides for 4h, in the presence of GolgiPlug and then stained for analysis of cytokine production and cytotoxicity. Results were compared by one-way ANOVA with Tukey correction. *P < 0.05; **P < 0.01; ***P < 0.001; ****P < 0.0001.

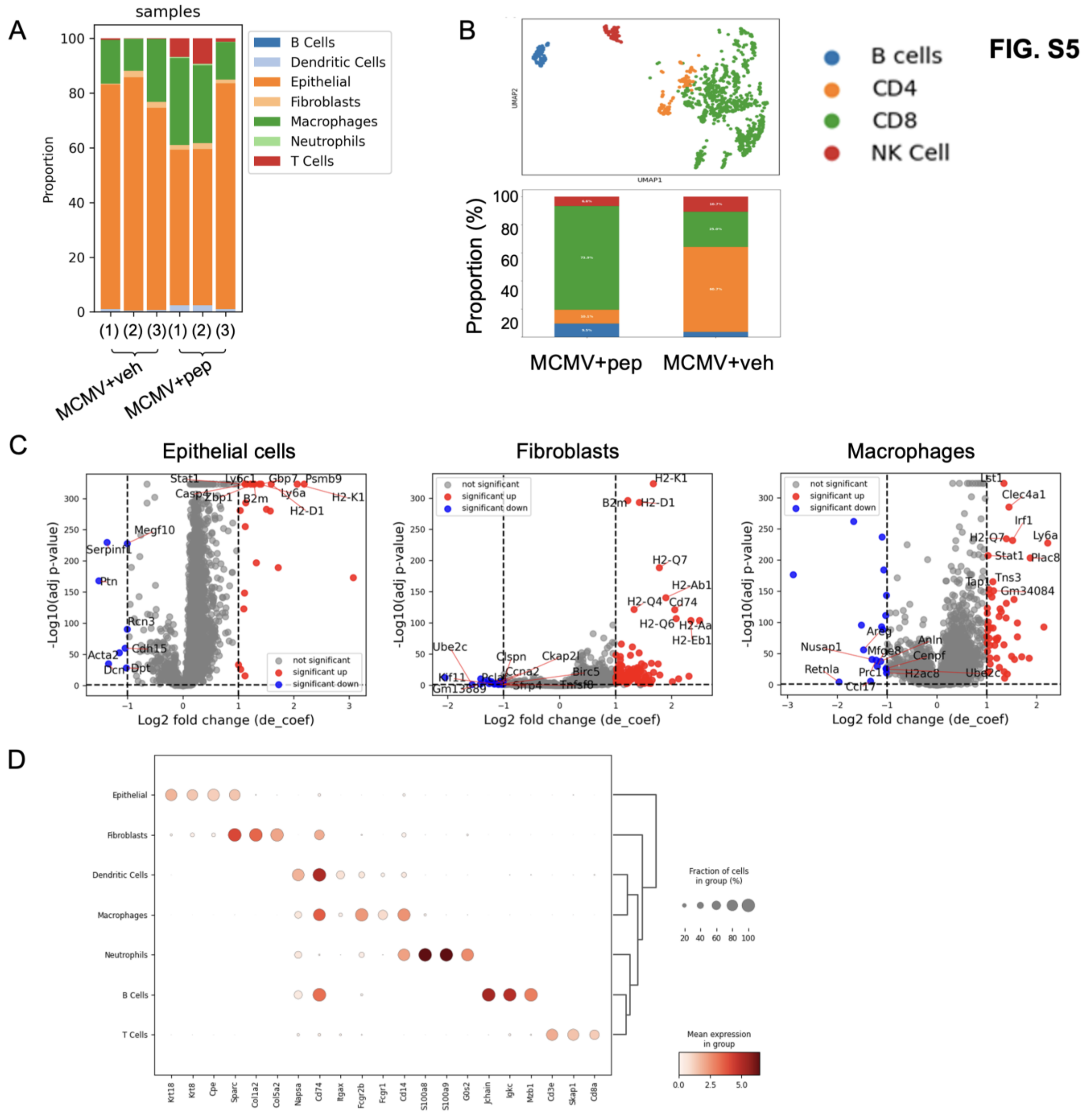


**Suppl Figure 5. Tumor infiltration by MCMV-specific T cell is accompanied by changes in the TME.** Additional analysis from the data shown in Fig 5. **A**, Cell type distribution per sample. **B**, UMAP sub-clustering of lymphocytes displaying B cells, T cells, and NK cells. **C**, Volcano-plots representing DEGs in epithelial cells, fibroblasts and macrophages. **D**, Markers used to annotate the cell types in Fig. 5.

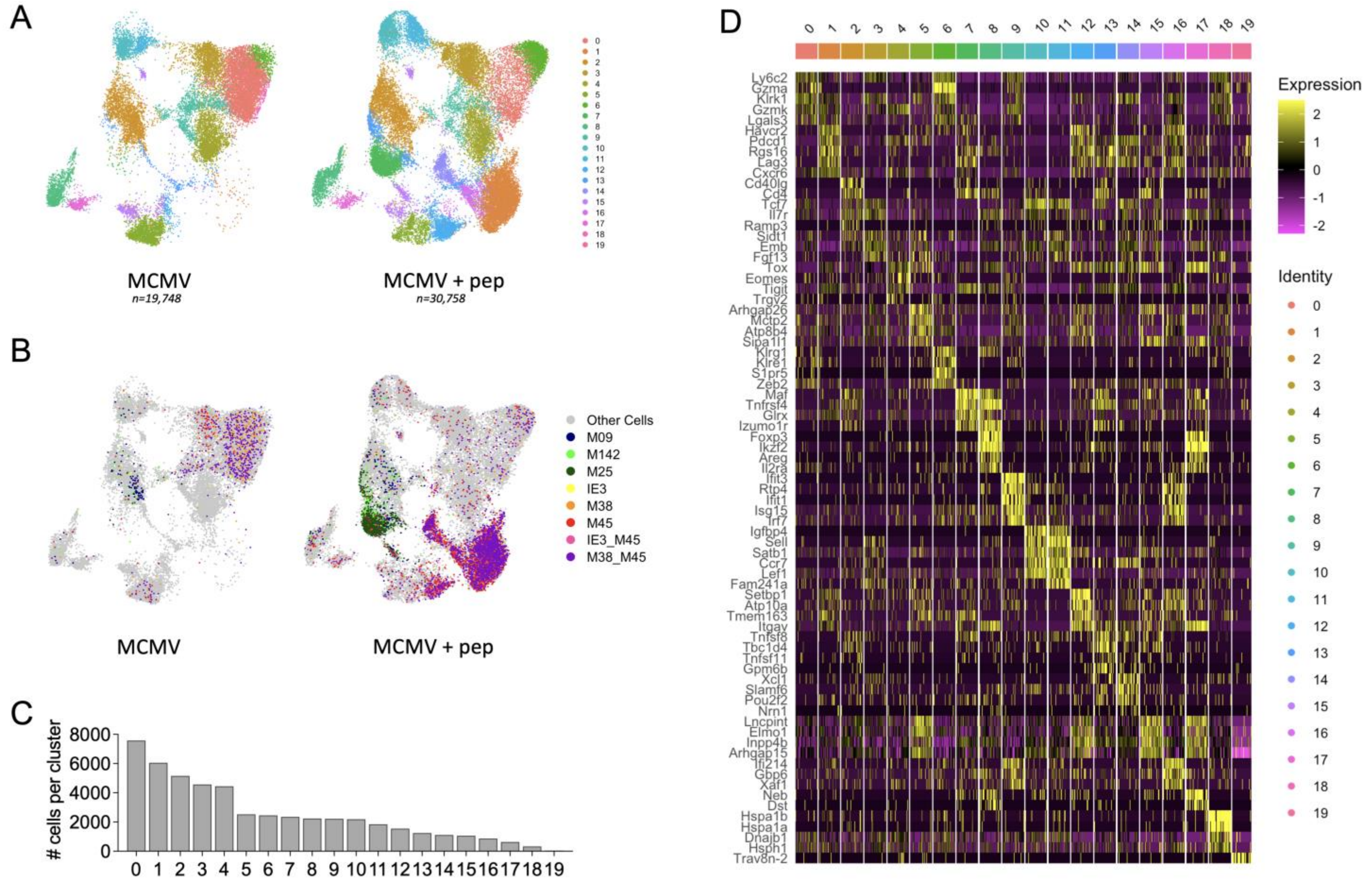


**Suppl Figure 6. Analysis of tumor infiltrating T cells.** Additional analysis from the data shown in Figure 6. **A**, T cell clustering and UMAP distribution split by sample conditions. **B**, Dimplot representing the detection of specific MCMV-associated matched CDR3 sequence for TCRβ. **C**, Absolute number of cells in each cluster. **D**, Top5 DEGs per cluster.

FIG. S7

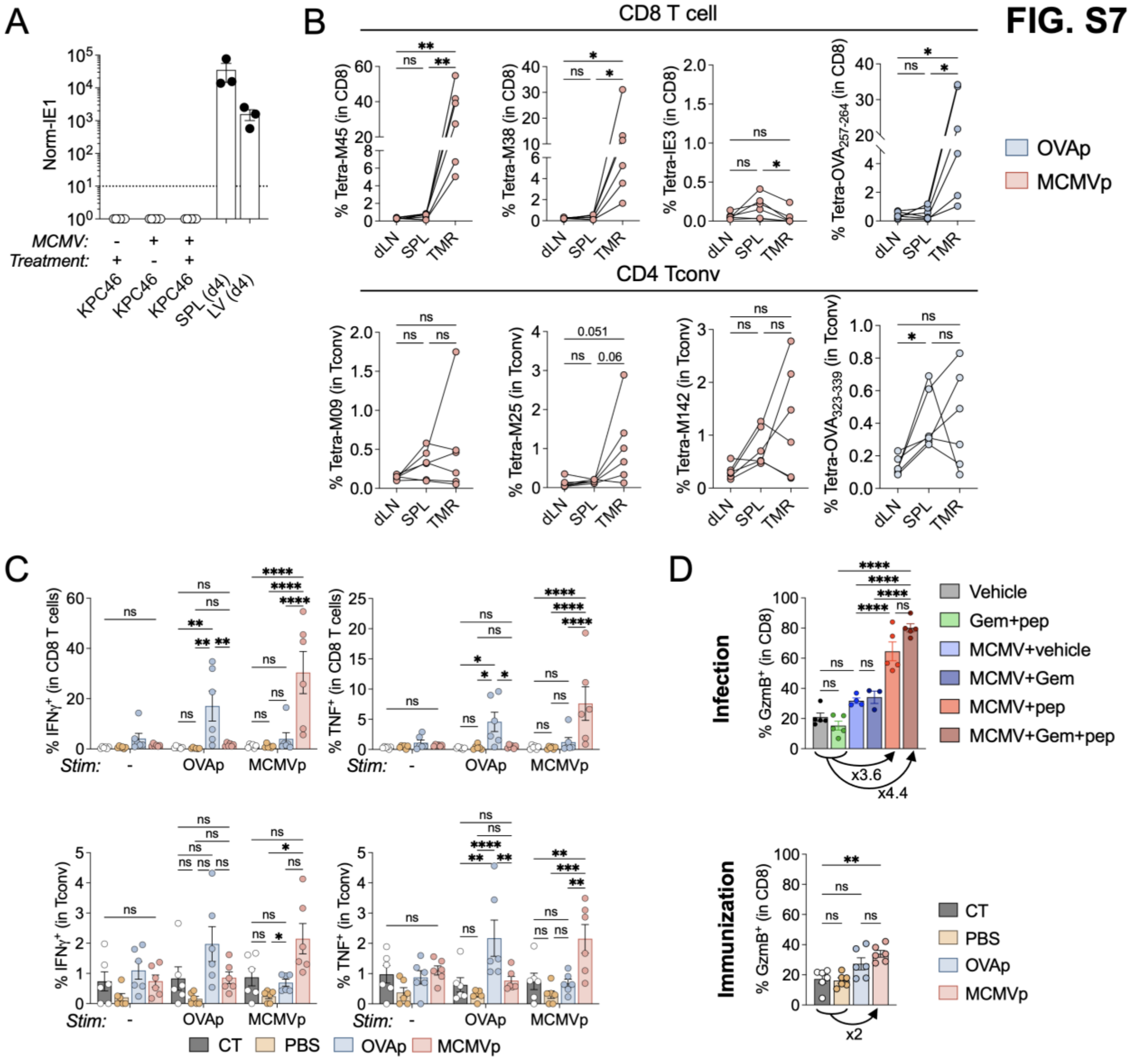


**Suppl Figure 7. Infection-induced T cells have more cytotoxic potential than immunization-induced T cells. A**, IE1 DNA qPCR normalized with actin, on KPC46 tumors harvested from uninfected or MCMV infected B6x129 F1 hybrid mice. Positive controls are spleen and liver from mice 4 dpi. **B**, Percentage quantification of MCMV and OVA tetramer binding T cells in MCMVp and SIINFEKL immunized mice from the draining lymph node, spleen and tumor. **C**, Purified CD45+ cells from tumors were incubated with the 6 MCMV peptides used for treatment, or the 2 OVAp for 4h, in the presence of GolgiPlug and IFNγ and TNF production was measured in CD8 and CD4 T conv cells. **D,** GzmB expression and fold increase in total CD8 T cells from the tumor, comparing control and treated groups after immunization (bottom) and infection (top, from experiment shown in Fig.4). Results were compared by one-way ANOVA with Tukey correction. *P < 0.05; **P < 0.01; ***P < 0.001; ****P < 0.0001.

| REAGENT | SOURCE | IDENTIFIER |
|---|---|---|
| Live-Dead eF506 | eBioscience | Cat# 65-0866-18 |
| Live-Dead Blue | Invitrogen | Cat# L34962 |
| RPE-streptavidin | Agilent | Cat# PJRS25-1 |
| APC-streptavidin | Agilent | Cat# PJ27S-1 |
| m09-biotinylated monomer (I-Ab) | NIH | GYLYIYPSAGNSFDL |
| m25-biotinylated monomer (I-Ab) | NIH | NHLYETPISATAMVI |
| m142-biotinylated monomer (I-Ab) | NIH | RSRYLTAAAVTAVLQ |
| m38-biotinylated monomer (H2-Kb) | NIH | SSPPMFRV |
| m45-biotinylated monomer (H2-Db) | NIH | HGIRNASFI |
| IE3-biotinylated monomer (H2-Kb) | NIH | RALEYKNL |
| OT-I-biotinylated monomer (H2-Kb) | NIH | SIINFEKL |
| OT-II-biotinylated monomer (I-Ab) (1) | NIH | HAAHAEINEA |
| OT-II-biotinylated monomer (I-Ab) (2) | NIH | AAHAEINEA |
| BUV496 anti-CD3e (17A2) | BD Biosciences | Cat# 751418; AB_2875417 |
| Pe-Cy7 anti-CD3e (17A2) | BioLegend | Cat# 100220; AB_1732057 |
| BUV395 anti-CD4 (GK1.5) | BD Biosciences | Cat# 563790; AB_2738426 |
| BUV805 anti-CD8 (53-6.7) | BD Biosciences | Cat# 612898; AB_2870186 |
| AF488 anti-Foxp3 (MF-14) | BioLegend | Cat# 126406; AB_1089113 |
| BV785 anti-CD44 (IM7) | BD Horizon | Cat# 563736; AB_2738395 |
| BV605 anti-CD44 (IM7) | BioLegend | Cat# 103047; AB_2562451 |
| APC-Cy7 anti-CD44 (IM7) | BioLegend | Cat# 103028; AB_830784 |
| APC-Cy7 anti-CD62L (MEL-14) | BioLegend | Cat# 104428; AB_830798 |
| BV570 anti-CD62L (MEL-14) | BioLegend | Cat# 104433; AB_10900262 |
| Pe-Cy5 anti-CD69 (H1.2F3) | BioLegend | Cat# 104510; AB_313112 |
| PercP-Cy5.5 anti-CD69 (H1.2F3) | BD Pharmingen | Cat# 561931; AB_394051 |
| BV785 anti-CD226 (DX11) | BD Biosciences | Cat# 744611; AB_2871596 |
| BV650 anti-LAG3 (C9B7W) | BioLegend | Cat# 125227; AB_2687209 |
| BV711 anti-PD1 (29F.1A12) | BioLegend | Cat# 135231; AB_2566158 |
| AF700 anti-TIM3 (#215008R) | R&D System | Cat# FAB1529RN; |
| AF700 anti-Ki67 (16A8) | BioLegend | Cat# 652419; AB_2564284 |
| Pe-Cy7 anti-GzmB (QA16A02) | BioLegend | Cat# 372214; AB_2728380 |
| eF450 anti-GzmA (GzA-3G8.5) | eBioscience | Cat# 48-5831-82; AB_2574079 |
| PE-Dazzle anti-Prf (S16009A) | BioLegend | Cat# 154316; AB_2922482 |
| APC anti-IFNg (XMG1.2) | BioLegend | Cat# 505810; AB_315403 |
| eF450 anti-IFNg (XMG1.2) | eBioscience | Cat# 48-7311-82; AB_1834366 |
| AF647 anti-TNF (MP6-XT22) | BD Pharmingen | Cat# 557730 |
| PE anti-TNF (MP6-XT22) | eBioscience | Cat# 12-7321-82; AB_466199 |
| BV605 anti-TNF (MP6-XT22) | BioLegend | Cat# 506329; AB_11123912 |

Table 1. Antibodies and reagent used for flow cytometry staining

*Whole Tumor scRNA-seq QC*

FASTQ files were processed using cellranger multi (cellranger-8.0.1) and aligned to cellranger-vdj-GRCM38. The resulting output files were then further filtered using cellbender (v0.3.2)(Fleming et al., 2023) using default settings. Afterward, droplets that had at least a probability of 0.5 of being cells were retained for further quality filtering. Cells were then filtered for outliers by counts using the median absolute deviation. Additionally, cells were also excluded if they had a mitochondrial RNA content of greater than 5%, or if they had less than 600 UMI counts. Doublets were then filtered out using scDblFinder (v 1.16.0) (Germain et al., 2022)

*Whole Tumor scRNA-seq annotation*
Batch correction was performed using scvi (v 1.1.2)(Gayoso et al., 2022). To determine ideal hyperparameters, a hyperparameter sweep was performed for 100 samples from the sample space. The model with the lowest validation loss was selected, and used for training the scvi model for 114 epochs. Using the computed latent space, nearest neighbors were calculated as well as a UMAP, using 30 nearest neighbors and a minimum distance of 0.1 for the UMAP. Cells were clustered using the Leiden algorithm implemented in scanpy (version 1.10.1) (Wolf et al., 2018), at a resolution of 0.6. Additionally, the igraph flavor was used, and it was run for 2 iterations. Clusters were then manually annotated using known marker genes (Fig. S7D)

*Whole Tumor scRNA Differential Gene Expression and GSEA*
For differential gene expression between conditions, we utilized the recently published memento package. We used the ht_1d_moments function, running for 40,000 bootstraps. All other arguments were left as default (Kim *et al.,* 2024). GSEA was then performed using ClusterProfiler (v 4.15.0.003) (Xu et al., 2024).

*Whole Tumor CellChat Analysis*
CellChat (v 2.1.2) (Jin et al., 2024) was used to perform CellChat analysis. The protocol described in the paper was generally used. Filtering was further performed to only include communications found in at least 10 cells and for pathways that were expressed in a minimum of 2 samples.

*TILs single-cell Transcriptome and TCR Repertoire*
Analysis of single-cell RNA sequencing (scRNA-seq) data were processed using the Cell Ranger v6.1.2 count pipeline for library mapping, followed by aggregation with the aggr pipeline. The aggregated data were imported into the R environment, and Seurat (v4.1.1) was utilized for quality control, normalization, and clustering. Cells expressing fewer than 200 or more than 2,500 genes, or with mitochondrial gene content exceeding 5%, were excluded. Genes detected in fewer than three cells were filtered out. The gene expression matrix was normalized and scaled prior to principal component analysis (PCA). Principal components were selected based on the elbow plot, and clustering was performed at a resolution of 0.8. Differentially expressed genes within clusters were identified using Seurat's FindAllMarkers function with the MAST statistical framework.T cell receptor (TCR) repertoire analysis was integrated with transcriptomic data using the SCrepertoire R package. The Seurat object was employed to organize both transcriptomic and clonotype metadata. TCR diversity was characterized through clonotype expansion analysis, V(D)J gene usage patterns, and CDR3 length distributions. Functions highlightClones() and combineExpression() were employed to map clonotype distributions onto UMAP projections. Antigen-specific TCR clonotypes for IE1, m38, and m45 were identified using the VDJ database (https://vdjdb.cdr3.net/) as well as an in-house TCR sequence library derived from tetramer-sorted cells. Cells identified with dual antigen specificity were manually reviewed to ensure proper annotation or reclassified as "other cells" when appropriate. Clonotype-specific transcriptional profiles were analyzed and visualized using UMAP embeddings, facilitating the identification of

functionally distinct T cell subsets. Statistical analyses were performed to correlate clonotype features with transcriptional states. Data visualization was conducted using custom scripts and ggplot2, leveraging the capabilities of both Seurat and SCrepertoire for integrative analysis.
